# Late Quaternary climatic impact on the woodland strawberry genome: a perennial herb’s tale

**DOI:** 10.1101/2024.10.09.617376

**Authors:** Tuomas Toivainen, J. Sakari Salonen, Jonathan Kirshner, Sergei Lembinen, Hanne De Kort, Annina Lyyski, Patrick P. Edger, Hrannar Smári Hilmarsson, Jón Hallsteinn Hallsson, Daniel J. Sargent, Klaus Olbricht, José F. Sánchez-Sevilla, Laura Jaakola, Johan A. Stenberg, Boris Duralija, Juozas Labokas, Henry Väre, Jarkko Salojärvi, Petri Auvinen, David Posé, Victor A. Albert, Timo Hytönen

## Abstract

Exploring a species’ paleohistory is crucial for understanding its responsiveness to climatic events, identifying drivers of adaptation, and developing effective biodiversity conservation strategies in the face of ongoing climate change. We analyzed 200 genomes of the perennial herb woodland strawberry (*Fragaria vesca* L.) from across Europe and investigated the population structure and demographic history of the species during past geoclimatic events. We found a clear division of populations into western and eastern genetic clusters, indicative of distinct glacial refugia and adaptations to variation in temperature seasonality. The eastern core populations were several times larger (defined as effective population size, N_E_) than populations in other regions, showed no evidence of inbreeding, and were resilient to several glacial maxima. However, we observed decreasing N_E_ and higher inbreeding in populations toward range edges, particularly in the north, where these individuals went through bottlenecks during glaciations. Population divergence suggested that western and eastern Europe were colonized from separate refugia in multiple waves during the Holocene, while the largest current populations from the northern Mediterranean to southern regions of the Nordic countries formed a connected population chain with gene flow between eastern core populations and western Europe, primarily occurring through Central Europe. Similar patterns of colonization and hybridization may have occurred during past interglacial periods, contributing to the present-day population structure of woodland strawberry. We suggest that the unprecedented resolution of the species’ climatic history across six glacial-interglacial cycles presented here holds the promise of transforming the general understanding of species paleohistory through geoclimatically tracing ancestral haplotypes.

## INTRODUCTION

Glacial-interglacial (GI) cycles during the Quaternary period, which lasted over the past 2.58 million years (Lisiecki and Raymo 2005), caused repeated distributional contractions and expansions of species (Hewitt 2000, Melles et al. 2012, Schiferl et al. 2023). Exploring these geoclimatic events that shaped the current spatial population genetic structures of species is crucial for understanding their responsiveness to future climatic events and drivers of adaptation, and for developing biodiversity conservation strategies. Before the advent of population genomics research, evidence for the influence of GI-cycles on species’ histories were obtained through analyses of dated paleoecological records, including fossil pollen found in sediments (Melles et al. 2012; Donders et al. 2021; Schiferl et al. 2023). Population genomics has become a powerful tool for investigating genetic diversity and demographic histories of species across past climatic events (hereafter, “climatic histories”) both in animals (Yim et al. 2014, Qiu et al. 2015, Liu et al. 2022, Wang et al. 2022, Kessler and Shafer 2024) and plants (Alonso-Blanco et al. 2016, Dong et al. 2023; Sun et al. 2020; Qiao et al. 2021, Milesi et al. 2023, Salojärvi et al. 2024). Recent studies in woody perennials, including wild grapevine (*Vitis sylvestris*) (Dong et al. 2023) and wild apple (*Malus sieversii*) (Sun et al. 2020), revealed significant declines in effective population sizes (N_E_) during the last glacial periods, i.e. the Last Glacial Maximum (LGM, 22-17 thousand years ago (ka), Lisiecki and Raymo 2005) and the Penultimate Glacial Period (PGP, 190-130 ka). However, several forest tree species with high N_E_ showed resilience to climatic events throughout the Quaternary period (Milesi et al. 2023).

Although perennial herbs comprise the largest fraction of plant species on Earth, with an increasing proportion of species towards colder environments (Chiarucci et al. 2017, Rice et al. 2019), their spatial genetic structures and climatic histories remain largely unknown. Studies of the perennial herbs alpine rockcress (*Arabis alpina* L.) and lyrate rock-cress (*Arabidopsis lyrata*) revealed strong population structures across Europe and stable climatic histories, and their northern populations displayed highly reduced genetic diversity, indicating colonization-associated founder effects (Laenen et al. 2018; Mattila et al. 2017). In the genus *Fragaria* L. (strawberry), creamy strawberry (*Fragaria viridis* Weston), the species with the highest determined N_E_ to date, showed greater resilience to the LGM climate than other species (Qiao et al., 2021).

In this study, we leverage whole genome sequencing data to investigate the population structure, current status, and climatic history of the perennial herb woodland strawberry (*F. vesca ssp. vesca* L.) across its European range. This major crop wild-relative of the Rosaceae family thrives in diverse habitats, including forests, meadows, and disturbed areas such as roadsides, and has a wide geographic distribution in Eurasia and as a non-native subspecies in eastern North America (Hilmarsson et al. 2017; POWO 2024). Woodland strawberry reproduces sexually through both outcrossing and self-fertilization, as well as asexually via above-ground stolons (Sargent et al. 2004). Flower buds are formed in short days in autumn, and develop flowers, ultimately fruits, the following growing season (Heide & Sonsteby 2007, Koskela et al. 2012). Birds and mammals eat the fruit and mediate seed dispersal (Herrera 1989, Heleno et al. 2010, López-Bao & González-Varo 2011). Here, we demonstrate the presence of distinct eastern and western genetic clusters in the European woodland strawberry and unveil the high-resolution climatic history of the species.

## RESULTS AND DISCUSSION

We gathered a collection of 200 woodland strawberry accessions covering the latitudinal range of the species in Europe (Fig. 1A; Hilmarsson et al. 2017) and sequenced their genomes to a mean depth of coverage of 15.3× using the Illumina platform. We identified 2.36 million biallelic single nucleotide polymorphisms (SNPs) and 343,311 biallelic small indels (≤40 base pairs (bp); 71.5% were shorter than 5bp) and used these data for population genomic analyses (Dataset 1).

**Figure 1.**
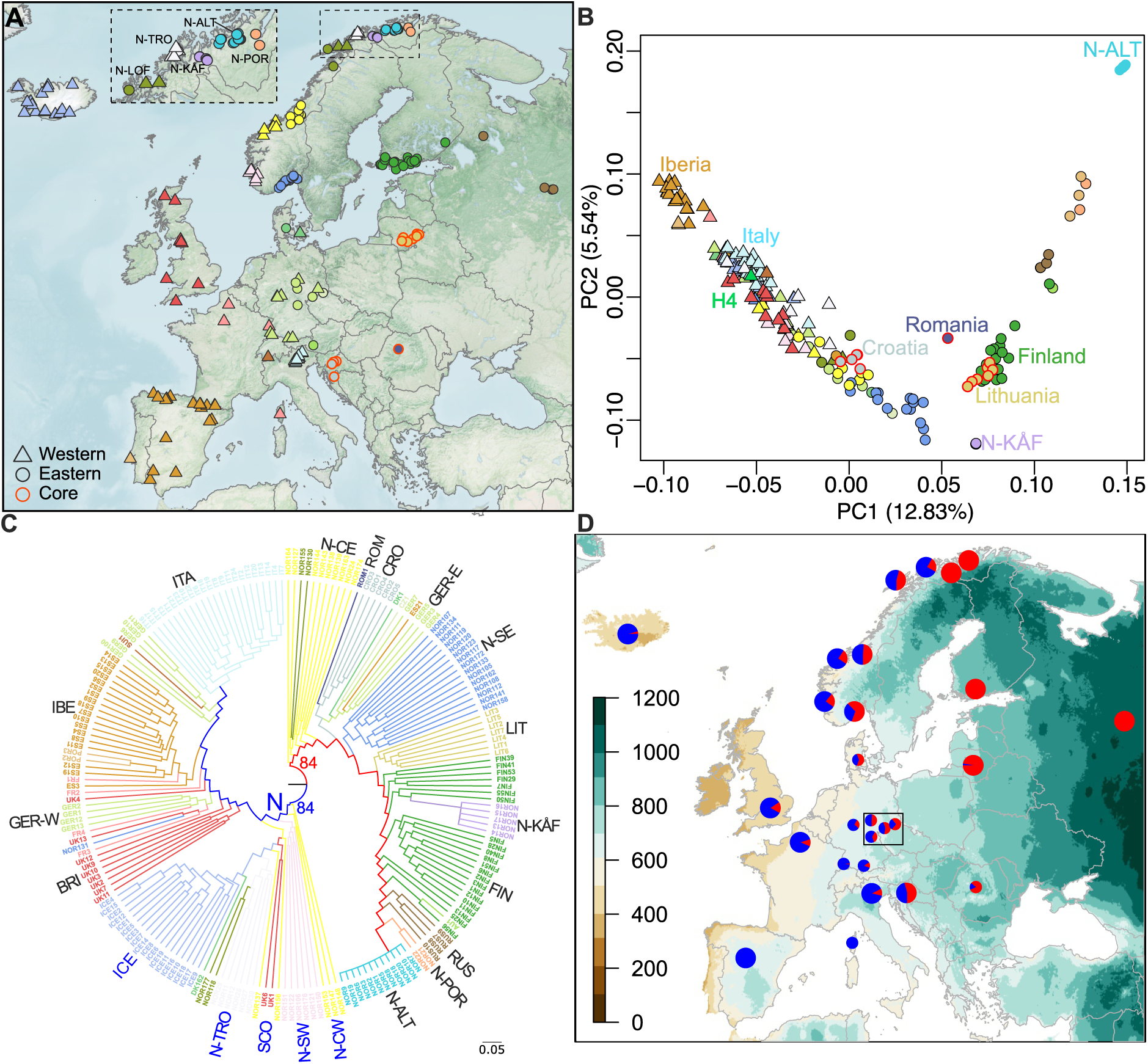
Collection sites and population structure of European woodland strawberry. A) Collection sites of sequenced woodland strawberry accessions. Western populations are indicated by triangles and eastern by circles; the division is based on the maximum likelihood phylogeny shown in C. Symbols with red outlines indicate core populations with large effective population sizes. B) Principal component (PC) analysis on the genomic data of woodland strawberry accessions shown in A using the same symbols. H4=Hawaii-4 reference accession (Edger et al. 2017). C) SNP phylogeny of European samples. The inner blue and red branches represent western and eastern samples, respectively. Bootstrap support values for these branches are shown. N indicates a northern sub-branch of the western branch, in which the names of sampled regions are colored blue for clarity. Outer branch colors correspond to colors in A and B. D) Average ancestry proportions derived from ADMIXTURE analysis (K=2), plotted for regional sample pools, from different countries/areas of countries or specific samples (large and small pies, respectively) on a background gradient map of temperature seasonality (BIO4; (standard deviation ×100). Blue=western ancestry, Red=eastern ancestry. German samples are shown with small symbols to highlight the gradient in the west-east admixture. The black rectangle shows the hybridization region. Abbreviations: BRI=Britain, CRO=Croatia, FIN=Finland, GER-W=Western Germany, GER-E=Eastern-Germany, IBE=Iberia, ICE=Iceland, ITA=Italy, LIT=Lithuania, N-ALT=Alta, N-CE=Central-Eastern Norway, N-CW=Central-Western Norway, N-KÅF=Kåfjord, N-LOF=Lofotes, N-POR=Porsanger, N-SE=Southeast Norway, N-SW=Southwest Norway, N-TRO=Tromsø, ROM=Romania, RUS=Russia, SCO=Scotland.

### Population structure suggests western and eastern glacial refugia in Europe

To explore the connection between population genetic structure and the geographic distribution of the woodland strawberry across Europe (Fig. 1A), we performed principal component, phylogenetic and admixture analyses. The first principal component (PC1), accounting for 12.9% of the variance, revealed two main branches that overlapped with two major clusters in maximum likelihood phylogenetic analysis of SNP data (Fig. 1B-C, S1B-C, S1E-F): western European samples from the Mediterranean to northwestern Norway spread along the left branch, while more easterly samples from Romania to northeastern Norway scattered along the right branch (Fig. 1B, fig. S1B-C). A Northeast-Southwest PCA cline is particularly visible in inverted form on the lefthand side of Fig. 1B. This strong geographic patterning likely reflects isolation by distance (IBD) phenomena (Wright, 1943, as exemplified by European humans; Novembre et al. 2008), wherein stepwise fixation of SNPs via random genetic drift in postglacially migrating populations emerged.

Specifically, the samples from the southern European peninsulas, Iberia (Spain and Portugal), Apennine (Italy), and the Balkans (Croatia), formed separated groups along the western branch, with the Croatian samples grouping closest to the eastern branch. The eastern populations from Kåfjord and Alta, closely located in adjacent fjords in northeastern Norway, were the most distinctly separated by the second PC (5.58%). These leading-edge populations also formed distinct groups in the SNP phylogeny and showed high genetic differentiation with an F_ST_ value of 0.77 (Fig. 1C, Table S1), suggesting strong bottlenecks during past colonization.

The geographic border between the western (N=112) and the eastern (N=87) European populations, as determined robustly by SNP phylogeny (Fig. 1C, Fig. S1D-F), extended from northern Norway through central Norway, southern Norway, and central Europe to Croatia in southern Europe (Fig. 1A). To explore the geographic distribution of western and eastern ancestries in greater detail, we conducted an ADMIXTURE-analysis (Alexander et al. 2009). Samples originating from near the border, for example in Croatia, Germany, and central Norway, displayed both eastern and western ancestries (K=2) in their genomes (Fig. 1D, Fig. S1G). In contrast, samples from Finland, Russia, Alta, and Kåfjord exclusively carried ancestries from the east, whereas Iberian samples were solely of western ancestry.

The border stretching from the Adriatic Sea to the Baltic Sea that separated western and eastern woodland strawberry also delineated the geographic ranges of two subspecies of the house mouse (*Mus musculus domesticus* vs. *M. m. musculus*) and the hedgehog (*Erinaceus europaeus* vs. *E. roumanicus*), as well as two admixture groups of silver birch (*Betula pendula*) and Chalk-hill blue butterfly (*Polyommatus coridon*) (He et al. 2012, Phifer-Rixey and Nachman 2015, Milesi et al. 2023, Schmitt and Zimmermann 2012). This demarcation suggests that western and eastern European populations of various organisms were isolated in distinct refugia during glacial periods, as suggested previously (Taberlet et al. 1998, Hewitt 2000). The border aligned with the primary hybridization zone in central Europe, where various species experienced secondary contact during postglacial range expansions (Hewitt 2000, Schmitt 2007). The fact that the populations followed the border between the temperate oceanic and humid continental climatic zones in western and eastern Europe (Beck et al. 2023) respectively, which differ in temperature seasonality (Fick and Hijmans 2017), similarly suggests that the gradient in temperature seasonality contributed to the differentiation of western and eastern genomes in the woodland strawberry (Fig. 1D, S2) and other species.

### Habitat fragmentation increases with latitude

To explore the present state of woodland strawberry populations, we studied the current N_E_ of each sample by analyzing their most recent coalescence rates (Schiffels and Durbin 2014, Wang et al. 2020), and assessing the extent of inbreeding by calculating inbreeding coefficients (F_IS_) within regional sample sets collected from different countries or areas of countries (see Materials & Methods). The effective population sizes of eastern European samples from Lithuania, Croatia and Romania were 5 to 10 times larger than the median N_E_ values in other regions (Fig. S3, Fig. S4). These samples were also located in the core of the PCA plot, hereafter termed “core populations” (Fig. 1A-B). The lowest N_E_ values were observed at the range edges specifically in the Iberian Peninsula, Iceland, and northern Norway (Fig. S3, Fig. S4). Further, strong negative latitudinal correlations in regional N_E_ values were found in both western and eastern Europe when the samples from southern and northwestern Iberia were excluded from the analysis (Fig. 2A and 2B). Aligning with N_E_, sample-specific nucleotide heterozygosity also decreased towards higher latitudes (Fig. S5A-B). However, regional nucleotide diversity (expected heterozygosity) did not correlate with latitude (Fig. S5C-D), suggesting that samples were increasingly differentiated towards the north due to habitat fragmentation and a higher rate of inbreeding.

**Figure 2.**
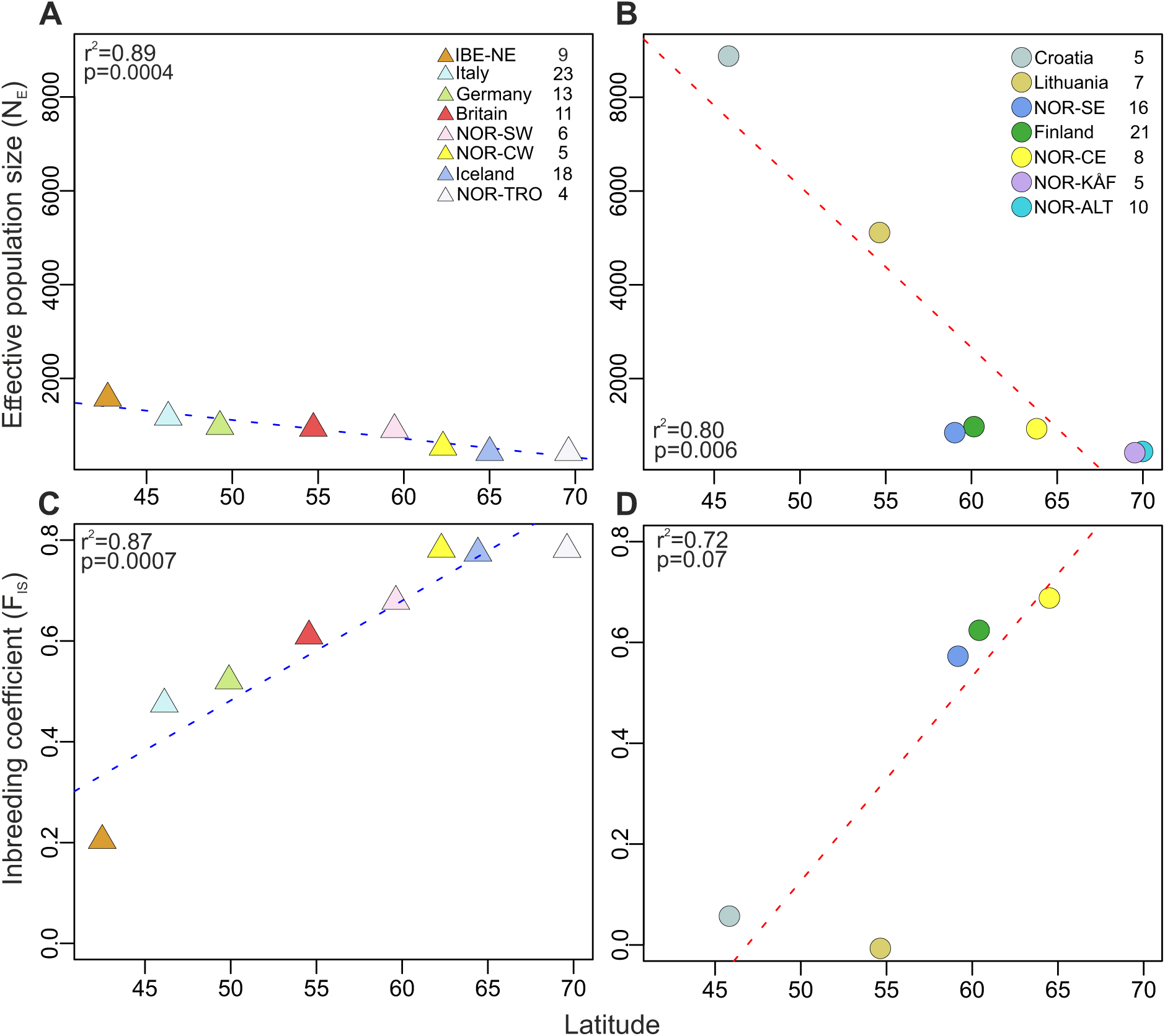
Latitudinal patterns of effective population sizes and inbreeding coefficients. A, B) Latitudinal correlations of median effective population sizes (N_E_) in western (A) and eastern (B) European samples. C, D) Latitudinal correlations of inbreeding coefficients (F_IS_) of regional sample sets from western (C) and eastern (D) Europe. The number of samples from each region is indicated in the legends after the name of each region. The coefficient of determination (r^2^) is derived from the Pearson correlation coefficient (r), with statistical significance indicated by the p-value.

Based on F_IS_ values, only woodland strawberry populations from Lithuania and Croatia were close to Hardy-Weinberg equilibrium with F_IS_ values close to 0 (Fig. 2D, S3B), indicating that populations in these regions exhibited panmictic characteristics with low levels of habitat fragmentation and inbreeding. In contrast, isolated populations in Alta and Kåfjord showed negative F_IS_ values (−0.12 and −0.22), suggesting an excess of heterozygous sites within samples, which could be related to asexual reproduction. However, because of very low nucleotide diversity and high genetic differentiation in these populations compared with other populations (Fig. S5D, Table S1), they were prone to analytical errors due to, e.g., poor read-mapping to the ‘Hawaii-4’ reference genome, which is closely related to Italian samples (Fig. 1B). Excluding Alta and Kåfjord, which were the only regions where F_IS_ did not correlate with N_E_ (Fig. S6), the correlation between N_E_ and F_IS_ was strongly negative and highly significant (r^2^=0.65, p < 2.2 × 10^-16^). When excluding extreme samples with potentially unreliable results or different modes of reproduction, a positive latitudinal correlation was found in F_IS_, especially in the western genetic cluster, ranging from 0.2 in northeastern Iberia to 0.8 in Iceland and Tromsø in northwestern Norway (Fig. 2C and 2D). Taken together, the strong latitudinal correlations in N_E_ and F_IS_ suggest strong founder effects in northern populations and increasing inbreeding towards the north, as expected after postglacial range expansions (Excoffier et al. 2009). Additionally, a decline in habitat quality towards the north may have contributed to shaping these correlations. Another study showed core-periphery effects specifically in animal-dispersed plant species, suggesting that this mode of dispersal, also used by woodland strawberry, is particularly sensitive to habitat fragmentation (De Kort et al. 2021).

### Southern refugia of woodland strawberry span four ice ages

To explore the demographic history of woodland strawberry, we analyzed population divergence and calculated time-dependent coalescence rates from whole, diploid genomes (Schiffels and Durbin 2014). We then applied an isolation-migration (IM) model to these rates (Wang et al. 2020), revealing changes in the historical effective population sizes and migration rates between populations. As in previous studies in other perennial herb species (Savolainen and Kuittinen 2011, Koch et al. 2006), we assumed a two-year generation time and the experimentally tested mutation rate of *Arabidopsis thaliana* (7.1*10^-9^ mutations per nucleotide per generation; Ossowski et al. 2010) in our demographic inference. We took advantage of published climatic data (Lisiecki and Raymo 2005) to support our demographic analyses, particularly regarding past glacial-interglacial cycles.

Given that southern European peninsulas (Iberia, Apennine, and the Balkans) have served as crucial refugia for plants and animals throughout the Pleistocene (Taberlet et al. 1998, Hewitt 2000), we investigated the timing of initial genomic divergence (M<0.999 and M<0.99) in populations currently inhabiting these areas (Figure S7A-B). Among the sample pairs among the three southern European peninsulas, and especially between Iberian and Romanian/Lithuanian populations, initial divergences consistently dated to 450 - 330 ka, and in some comparisons to over 600 ka (Fig. S7A, B). This period spans the transition from marine isotope stage 11 (MIS 11) to the MIS 10 glacial period, when migration between samples from Iberian, Apennine and Balkan peninsulas often ceased (Table S3, see below). These findings suggest that distinct woodland strawberry evolutionary lineages were established in all three peninsulas at least 330 ka, spanning four ice ages. Drawing parallels with other species, Hewitt (2000) used mitochondrial sequences to estimate that *Ursus arctos* (brown bear) and *Chorthippus parallelus* (grasshopper) likely sought refugia in these areas around the same time.

### Core and peripheral populations show contrasting demographic patterns

Our analyses of historical N_E_ and migration rates revealed two common contrasting patterns that aligned with the chronological timing of serial MIS (Text S1, S2): a “core pattern” showing stable N_E_ and migration rate across multiple GI-cycles until the LGM (Fig. 3A, 3C, S10, S11), and a “peripheral pattern” characterized by its susceptibility to bottlenecks and isolation events during glacial maxima and rehybridizations during following interglacial periods (Fig. 3B, 3D, S12, S13). The stable core pattern was most frequently revealed when haplotypes were drawn from two eastern core populations with high N_E_ from Croatia, Romania, or Lithuania, but other haplotype combinations rarely (*p* = 1.22*10^-6^) showed this pattern (Table S3, Fig. S17B). The climate sensitive peripheral pattern was typically found in sample pairs from different peninsulas but also between northern and southern samples (Fig. 17B-C). These patterns broadly resembled the eastern (core) and the western (peripheral) demographic histories of wild grapevine (Dong et al. 2023).

**Figure 3.**
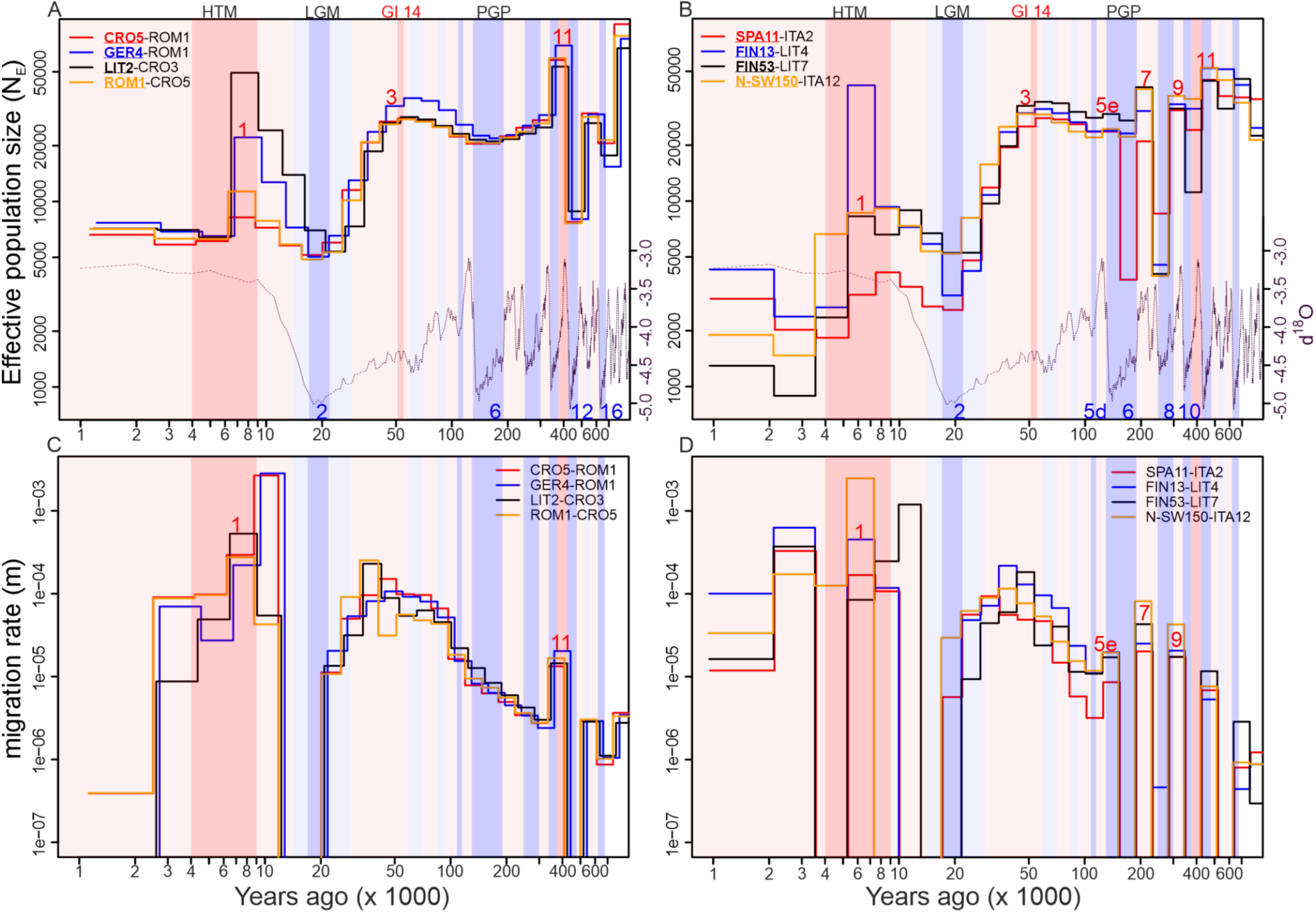
Demographic history of European woodland strawberry. A, B) Effective population size (N_E_) in the core (A) and the peripheral pattern (B) through time using MSMC-IM. The trajectory of the first sample from each sample pair is shown. C, D) Historical migration rate (m) in the core (C) and the peripheral (D) patterns. X-axis values represent thousands of years. Glacial and interglacial periods are shown with background colors, and blue and red numbers indicate specific marine isotope stages (MIS) and substages (a letter after the number). Curves at the bottom of A & B represent inverse benthic δ18O records from Lisiecki and Raymo (2005) and are used as a proxy for historical temperature. HTM=Holocene thermal maximum (9-4 ka), LGM=Last glacial maximum (22-17 ka, based on Lisiecki and Raymo 2005), GI14=Greenland interstadial 14 (55-51 ka) and PGP=Penultimate Glacial Period (190-130 ka).

The core pattern found in 7.6% of MSMC-IM runs demonstrated a history spanning 650 ka (Fig. 3A, 3C, S10, S11), with two bottlenecks in the deep past, the first possibly during the MIS 16 and the second during the MIS 12 glacial period (Fig. 3A and Fig. S10A-C), with the MIS 12 indicated as the most severe of all Pleistocene glacial periods in terms of ice extent and temperature (Hughes and Gibbard 2018). The MIS 12 glaciation resulted in particularly strong bottlenecks (Fig. 3A, S10A-C, S11A-C), and complete cessation of migration (Fig. 3C, S10D-F, S11D-F, S14A, S16), as also observed from ancestral tree pollen samples from Lake Tenaghi Philippon (Tzedakis et al. 2006) and Lake Ohrid in the Balkans (Sadori et al. 2016, Kousis et al. 2018, Koutsodendris et al. 2019). After the MIS 12 bottleneck, N_E_ values increased up to ten-fold (Fig. 3A, Fig. S10A-C, S11A-C), possibly during the unusually long MIS 11c super-interglacial period (426-396 ka), when the largest temperature increase during the late Quaternary was observed (Kousis et al. 2008, Melles et al. 2012, Marino et al. 2018). Following a noticeable decrease of N_E_ after the MIS 11 interglacial period, N_E_ was relatively stable (> 20,000) throughout GI-cycles, showing only slight decrease towards the PGP that was identified as a stronger glacial period compared to the earlier MIS 10 and MIS 8 glaciations (Fig. 3A) (Lisiecki and Raymo 2005). This observation, together with stable migration rates observed between the core populations across GI-cycles (Fig. 3C, fig. S10D-F, S11D-F), suggests ancestral haplotypes with the core pattern originated from a southern refugium that remained intact during glacial periods. The frequent emergence of the core pattern from analyses of Croatian and Romanian haplotype pairs (Fig. S17C) suggests that a stable refugium for woodland strawberry may have existed in the Balkan Peninsula, a region widely considered to have served as a major refugium for various species during the Pleistocene (Hewitt 2000; Taberlet et al. 1998).

Peripheral patterns, showing a bottleneck and/or cessation of migration between MIS 10 and the LGM, were found in more than 55% of MSMC-IM runs (Fig. S17A, Text S1). The history of the peripheral patterns extended up to 370 ka, corresponding to the MIS 10 glacial, with sufficient resolution for chronological marine isotope stages (Fig. 3B, 3D, S12, S13). A strong peripheral pattern, found in 19% of runs (Peripheral-1; Text S1, Fig. S17A), showed high sensitivity to consecutive marine isotope stages, with N_E_ fluctuations and migration rate patterns associated with GI-cycles across MIS 10 - MIS 5 (Fig. 3B, 3D, S12, S13). In different samples, strong bottlenecks were detected during either MIS 8 or the PGP (Fig. 3B, fig. S12A-C, S13A-B), and moderate bottlenecks were also associated with MIS 10 (Fig. S12A-C).

Consistent with the bottlenecks, migration most often completely ceased during these three glaciations (Fig. 3D, S12D-F, S13C-D, S14B, S15), whereas interglacial periods (MIS 9 and MIS 7) led to higher migration rates compared to the core pattern, indicating cycles of divergence and hybridization between populations; a phenomenon not detected in the core pattern of history (Fig. 3C, S10D-F, S11D-F).

### LGM caused a strong bottleneck in all European woodland strawberry populations

Following the PGP, N_E_ values increased in all samples towards the MIS 3 interstadial (59-29ka), reaching their maximum around the Greenland Interstadial 14 (GI14, 55-51ka), which was characterized by a climate with nearly present-day summer temperatures across Europe (Helmens et al. 2007, Meerbeeck et al. 2011, Sirocko et al. 2016). After GI14, N_E_ values began to decline at an accelerating pace in all samples, reaching a minimum at the onset of the LGM (Fig. 3A-B, fig. S10A-C, S11A-C, S12A-C, S13A-B). Concurrently, gene flow between populations abruptly ceased (Fig. 3C and 3D, fig. S10D-F, S11D-F, S12D-F, S13C-D), indicative of harsh conditions impeding migration. Unlike the three previous glacial periods, core populations also underwent significant bottlenecks during the LGM, accompanied by a cessation of gene flow between them. The most pronounced declines of N_E_ during MIS 2 were observed in populations currently at the range edges, including samples from, Alta (Fig. S18C, *p* = 5.05E-5), Kåfjord (Fig. S18A, p = 2.96E-5), Finland (*p* =1.9E-5), and the Iberian Peninsula (*p* = 3.87E-5). These declines were notably significant when compared to the N_E_ in Croatia or e.g. Italy during MIS 2 (Table S4).

Around the onset of MIS 1 at 14 ka and the following transition to the Holocene at 11.7 ka, N_E_ values started to increase, reaching their apex (Fig. 3A and 3B, fig. S10A-C, S11A-C, S12A-C, S13A-B) during the first half of the Holocene Thermal Maximum (HTM, ca. 9-4 ka, Salonen et al. 2011, Renssen et al. 2012, Mauri et al. 2015), which has been broadly recorded, especially in central and northern European paleoclimatic data (Mauri et al. 2014, 2015). A notable exception occurred in Lithuania, where effective population sizes seemed to surge dramatically, increasing even ten-fold (N_E_=50-70k) compared to the LGM (Fig. 3A, fig. S10A, S11A, Table S5). In southern European populations, the observed increase in N_E_ values was more modest, ranging from 1.5 to 3-fold (Fig S10A-C, Fig. S11A-C, Table S5), which aligns well with European paleoclimatic data (Mauri et al. 2014, 2015). Following peak N_E_ during the HTM, population sizes began to decrease, with stronger declines towards northern latitudes (Fig 3A, 3B, S10A-C, S11A-C, S12A-C, S13A-B, S18A, S18C, S19, Table S5). In these regions, climatic events can amplify temperature changes several-fold compared to the global average, as observed over the past 40 years (Rantanen et al. 2022). This could potentially lead to strong founder effects. Current N_E_ values are significantly lower compared to those of earlier interglacial periods (Fig. 3A and 3B, S10A-C, S11A-C, S12A-C, S13A-B), a trend also observed earlier in woodland strawberry and other plant species (Durvasula et al. 2017, Mattila et al. 2017, Qiao et al. 2021). This indicates a higher degree of habitat fragmentation during the late Holocene compared to earlier interglacial periods, coinciding with deforestation that started in Europe 5 ka (Marquer et al. 2014).

### Northern Europe was colonized from both sides of the continent in several waves

Divergence analyses shed new light on the postglacial migration of woodland strawberry that has led to its current spatial population genetic structure. Although there were early signals of divergence between putative refugia (Fig. S7), most of the genome remained undiverged (median cumulative migration probability, M>0.5) until the LGM, when gene flow ceased between populations (Fig. 3C, D, S10D-F, S11D-F, S12D-F, S13C-D), and splits between several western (Iberian) and eastern (Romanian or Lithuanian) samples were observed to have occurred (Fig. 4A, S20C-D, Table S3). These splits coincided with the separation of predicted ecological niches for western and eastern ecotypes of wild grapevine during the LGM (Dong et al. 2023), further supporting the impact of the LGM on population divergence. Additionally, the decline in their effective population sizes towards the LGM certainly contributed to the separation of western and eastern ancestries in both woodland strawberry and grapevine (Fig. 3A-B; Dong et al. 2023). Further analyses showed that the median split time between all western and eastern sample pairs (9819 ya) was significantly earlier than among western (5632 ya, *p*=1.02*10^-12^) and eastern (5723 ya, *p*=2.15*10^-12^) population pairs (Fig. S20B, Table S6), supporting the idea that western and eastern Europe were mainly colonized by western (i.e. Iberian or Italian) and eastern (i.e. Balkan) ancestry respectively, during the Holocene (Fig. 4A). As indicated by the cessation of gene flow during glacial periods, repeated splits of western and eastern populations likely occurred when plants retreated to western and eastern refugia during glacial maxima, but they were hidden by mixing of genomes across Europe during following GI-cycles (Fig. 3C, 3D, S12D-F, S13C-D).

**Figure 4.**
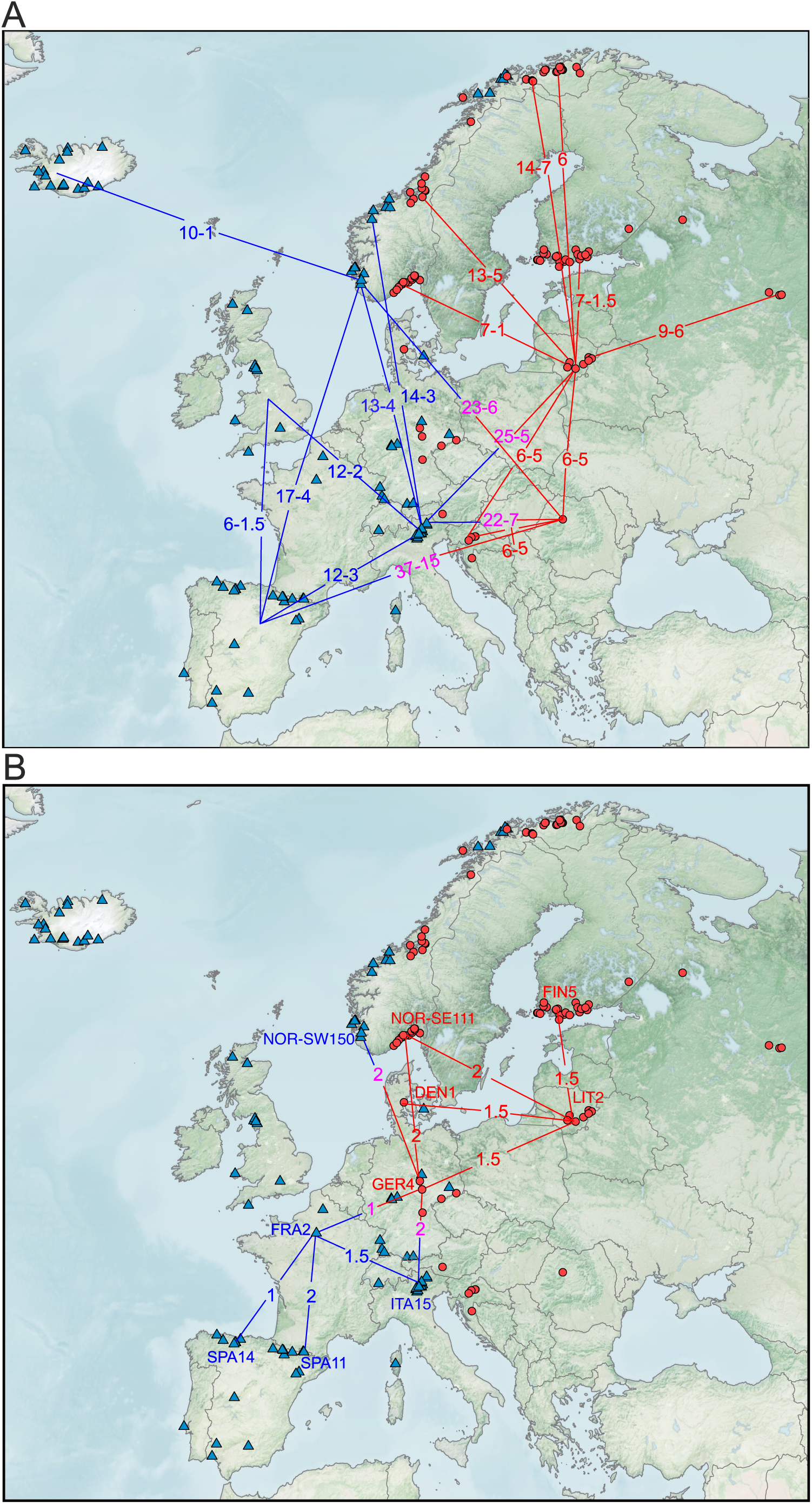
Divergence in European woodland strawberries. A) Ranges of split times in thousands of years between the regional samples during the postglacial northward migration of woodland strawberry. B) Split times between the largest regional populations from northern Iberia to southern Scandinavia. Blue and red lines/numbers indicate divergence within the western and eastern genetic cluster, respectively, while magenta numbers show the split time (ka) between western and eastern samples.

In eastern Europe, the Lithuanian population split significantly later from other eastern core populations, i.e., from Croatia (*p*=7.3 × 10^-04^) and Romania (*p*=3.2 × 10^-05^), than from Italy (Fig. S20C, Table S6), supporting the eastern colonization route from the Balkans. Peripheral populations from Alta, Kåfjord and Russia, and some samples from Finland and southeastern Norway diverged from Lithuanian samples during the first half of the Holocene, mostly during the early HTM (9-6 ka) (Fig. 4A, S20A-C), which saw amplified warming in the European Arctic (Salonen et al. 2011; Huntley et al. 2013; Väliranta et al. 2015), while the rest of the eastern samples split during the late HTM (6-4 ka) (Fig. S20A). The early split of Alta and Kåfjord samples from Lithuanian samples was followed by a strong bottleneck with a transient N_E_ decline of 98.3% and 96.0% compared to peak N_E_ during the HTM, indicating that eastern woodland strawberry ancestry reached the Arctic region during that time.

In western Europe, Britain was colonized from the western ancestry, since the split times of British samples from eastern populations (Croatia, Lithuania or Romania) were significantly more distant (median p-value=0.0036 for six comparisons) than from western populations (Iberia and Italy) (Table S6, Fig. S20C). However, the split times between British and Iberian samples were generally more recent than their divergence from Italian samples. This suggests a major contribution of Iberian ancestry to the colonization of Britain, a finding further supported by admixture analysis (Fig. S1G). Germany, situated on the border between eastern and western genetic clusters, was primarily colonized from Italy, as indicated by significantly closer split times from Italy compared with Iberian (p=0.004), Croatian (p=0.014), Lithuanian (p=0.034), or Romanian (p=0.001) samples (Table S6, Fig. S20C).

Norwegian coastal regions (NOR-SW, NOR-CW, NOR-CE) were likely colonized from both sides of Europe, since there were no significant differences in split times from western (Iberia or Italy) and eastern (Lithuania or Romania) ancestries (Table S6, Fig. S20C). While these samples mostly split during the HTM, there were several northwestern individuals from Scotland (UK1 and UK6), SW-Norway, CW-Norway, Iceland and Tromsø that split much earlier, certainly before the Holocene (Table S3, Fig. S20C). Their effective population sizes were usually much lower already by MIS 2 compared with populations that diverged during the HTM. Additionally, these samples were situated in the northern sub-branch of the western branch in the SNP phylogenetic tree (Fig. 1C). Together, these findings suggested that substantial portions of these particular genomes originated from a northwestern glacial refugium, similar to what has been suggested for *A. lyrata* (Mattila et al. 2017). Finally, several Icelandic subpopulations diverged from southwestern Norway only 2-1,3 ka (Fig. 4A, Fig. S20C, Table S3), indicating that the most recent colonization of woodland strawberry in Europe occurred in Iceland.

Further analysis of samples with the largest current N_E_ from Iberia (SPA14 and SPA11), France (FRA2), Italy (e.g. ITA15 and ITA2), Germany (GER4), Lithuania (LIT1 and LIT2), southeastern Norway (NOR-SE111 and NOR-SE120) and Finland (FIN5 and FIN13) revealed recent split times from their closest neighbors, comparable to split times between Lithuanian subpopulations (Fig. 4B, Table S3). This implies that the largest populations, of which several were hybrids, formed a connected population chain stretching from northern Spain and Italy to southern parts of the Nordic countries, with gene flow between eastern and western Europe primarily occurring through Germany (Fig. 4B). The Croatian and Romanian populations are not part of this chain, however, and should history repeat itself during future glacial periods, the Balkan peninsula may serve as a haven for the genetic diversity of the species.

## CONCLUSIONS

We demonstrated the division of the perennial herb woodland strawberry into western and eastern genetic clusters across Europe, with the largest (high N_E_) core populations occurring in (south)eastern Europe and the smallest populations in the range edges, particularly in the north. Eastern and western glacial refugia and different climatic conditions, such as differences in temperature seasonality, may have had a role in defining this demarcation in woodland strawberry and other species during the Quaternary. Consistent with ecological theory (Eckert et al. 2008), we found highly different demographic patterns between the core and other (peripheral) populations. Peripheral populations were susceptible to bottlenecks and isolation, while ancestral populations in core regions formed a shared macro-refugium that persisted throughout multiple Pleistocene glacial cycles. This macro-refugium served as a crucial reservoir of genetic diversity, evident from the rehybridization between core and peripheral populations during interglacial range expansions.

More evidence for multiple refugia came from divergence analyses that showed the colonization of northern Europe from both sides of the continent in multiple waves during the Holocene. We also found increasing homozygosity and decreasing N_E_ towards the north as evidence of founder effects and more fragmented and isolated habitats in the harsher northern environment. However, the largest populations from northern Spain and Italy to the southern regions of the Nordic countries formed an interconnected population chain with continuous migration. Moreover, gene flow between eastern and western Europe seemed to occur primarily through Germany, near the hybridization zone common for several species. We propose that similar colonization and hybridization processes occurred repeatedly during multiple interglacial periods, shaping the current spatial genetic structure of European woodland strawberries.

Finally, our demographic inference using MSMC-IM provided a new level of resolution on the climatic history of the species. We demonstrated fluctuating N_E_ and migration rate patterns across six glacial and six interglacial periods over the last 600 ka, with the migration rate providing higher resolution in detecting glacial maxima compared to the traditionally used N_E_. In principle, the capacity to trace the genomic locations of ancestral haplotypes from distinct historical periods could lead to groundbreaking, general insights into the evolution and climatic adaptation of plant species.

## MATERIAL AND METHODS

### Samples, DNA extraction and sequencing

Wild accessions of woodland strawberry were collected as clones or seeds, or obtained as clones from existing collections at Hansabred, Germany and IFAPA, Spain. Single seedlings were raised from seeds collected from each location, and together with clonal plant materials, the collection (Figure 1A, Table S7), covering majority of the European distribution of the species (Hilmarsson et al. 2017), was maintained in a greenhouse at the University of Helsinki. For DNA extraction, young, folded leaves were collected and frozen in liquid nitrogen followed by DNA extraction using CTAB-protocol (Doyle & Doyle 1990). Sequencing libraries were constructed using Nextera DNA Flex (now DNA prep) protocol according to manufacturer’s instructions (Illumina). Sequencing was performed with an Illumina NextSeq 500 machine using a paired end sequencing runs (170 bp and 140 bp or 151 bp and 151 bp).

### Trimming, mapping and initial variant calling

Raw Illumina reads were trimmed and paired with Cutadapt (DOI:10.14806/ej. 17.1. 200) using -q 25 and -m 30 parameters. Trimmed reads were mapped with BWA-MEM (Li 2013) against the *F. vesca* ‘Hawaii-4’ reference genome v.4.0 (Edger et al. 2017) with -M option. Duplications were marked with the Picard tool MarkDuplicates. GATK v 3.7. HaplotypeCaller-function was used to call variants in single samples, and then, variants were jointly called across all samples using the GenotypeGVCFs-function. Indels were filtered with following parameters: QD < 2.0 || FS > 200.0 || SOR > 10.0 || InbreedingCoeff < −0.8 || ReadPosRankSum < −20.0, and SNPs: QD < 2.0 || FS > 60.0 || SOR > 4.0 || MQ < 40.0 || MQRankSum < −12.5 || ReadPosRankSum < −8.0.

### SNP panel

Following use of the GATK pipeline, SNPs and indels closer than 10bp from indels were removed with bcftools (Li 2011), which resulted in 3,479,750 variants (2,898,156 SNPs and 581,594 indels) across samples. On average, 5.3% of missing data per polymorphic site was found. Several additional filtering steps were performed to remove low quality SNPs using vcftools (Danecek et al. 2011) before imputation and phasing. Initially, we included only those sites that had a maximum of 10% missing data and a mean coverage of less than 29, approximately twice the average coverage. Excessively heterozygous sites were removed from all samples if they deviated from Hardy-Weinberg equilibrium (p<0.01) in any regional set of samples. Excessively homozygotic sites were not removed because of the self-compatible nature of *F. vesca*. After additional filtering, 2,720,726 variants remained. Imputation was conducted using BEAGLE v4.1 (Browning et al. 2014) with default settings. Phasing was conducted for all samples simultaneously using SHAPEIT v2 software (Delaneau et al. 2013). Phase informative reads (PIR) were utilized in phasing with the extractPIRs-tool included in the SHAPEIT-software. The following parameters were used in phasing: -rho 0.001 --states-random 200 --window 0.5 --burn 10 --prune 10 --main 50. Variants were annotated with SnpEff-software (Cingolani et al. 2012) prior to extracting 4-fold degenerate sites from synonymous sites.

### Population structure

Principal component analysis (PCA) was conducted with SNPrelate software (Zeng et al. 2012) using polymorphic 4-fold degenerate sites (39,410 SNPs), nonsynonymous sites (79,727 SNPs), and genome-wide using linkage disequilibrium pruned sites (R2<0.2 in 20kb region, 62,857 SNPs) with a minor allele frequency greater than 0.01. ADMIXTURE software (Alexander et al. 2009) was used to determine admixture proportions of genetic clusters for each sample. Cross-validation error was the lowest at K=8 (Fig. S1), which was selected to represent population structure in the admixture data set. To visualize eastern and western clusters across accessions, K=2 was used. A phylogenetic tree using the same sites and based maximum likelihood was constructed with IQtree (Minh et al. 2020) and the general time reversible (GTR) model of molecular evolution and Lewis ascertainment bias correction (Lewis 2001) for noncontiguous SNP data. Otherwise, default parameters were employed, including 1000 bootstrap replicates. Genetic differentiation between regions was estimated by weighted F_ST_ (Weir and Cockerheim 1984) for regional pairs of samples (N=5).

### Inbreeding coefficients

Inbreeding coefficients (F_IS_) were calculated with vcftools (Danesec et al. 2011) for samples in each region or country (Figure 2, Fig. S3B, Table S2) when at least four samples were available from the same geographic region. Note that our particular sampling biases results to some extent. We do not have fully population-level samples; instead we sampled one individual per population from several separate populations within regions. Thus, we have population structure within the regions (except in Croatia and Lithuania), which increases a proportion of homozygous genotypes within regions, a phenomenon also called the Wahlund effect (Wahlund and Sten 1928). Especially samples from Iberia, Germany and Britain have been collected from large geographic areas around the countries. Despite suboptimal sampling, the observed high correlation (r^2^ = 0.64) between N_E_ and F_IS_ across samples suggests that our F_IS_ estimates are broadly reliable. Two samples from Iberia (ES21 and ES3) and Tromso (NOR30 and NOR28) were excluded from analyses based on highly negative F_IS_ values compared to other regional values, probably caused by recent hybridization (Fig. S1G). Median F_IS_ and sample latitudes were calculated from each region to explore correlation between F_IS_ and latitude across Europe both in western and eastern European samples (based on SNP-phylogeny).

### Inferring demographic history of populations

Demographic history of populations was inferred from statistically estimated haplotypes. Multiple Sequentially Markovian Coalescent (MSMC)-based method MSMC2 (Schiffels and Durbin 2014) was used to estimate coalescence rates within and across pairs of populations. To obtain a time-dependent estimate of migration rate and effective population sizes for each population pair, a continuous Isolation-Migration model was fitted to coalescence rates using MSMC-IM-software (Wang et al. 2020). To ensure the highest possible quality of the SNP data for coalescence analyses, SNPs were post-filtered with recommended additional filters for each sample (bam-files) and universally for the genome. Specifically, the Bamcaller.py script (https://github.com/stschiff/msmc-tools/blob/master/bamCaller.py) was used to produce additional sample-specific masks for low quality SNPs potentially produced by the pipeline. Only sites with a base quality above 20 and read mapping quality of Q>20 were retained. Moreover, sites with unusually high (> 2 x sample mean depth) coverage were masked. In addition, Heng Li’s SNPable software (https://lh3lh3.users.sourceforge.net/snpable.shtml) was used to produce universal masks for 35bp regions (35mers) that were not found uniquely in the *F. vesca* genome (Edger at el. 2017).

First, the MSMC2 was ran using default parameters with 20 iterations and the -s option to skip sites with ambiguous phasing to uncover within and cross-population coalescence rates between sample pairs by comparing all haplotype pairs (0-2,0-3,1-2,1-3) between samples.

Then MSMC2-results were analyzed with the MSMC-IM software using default parameters and recommended beta values for regularization of gene flow (1e-8) and N_E_ (1e-6). Several ancestral N_E_ values (30 000, 100 000 and 1 000 000) were tested for different sample pairs, but chose the default ancestral N_E_ (15 000), since it clearly had the highest sensitivity to detect the demographic events in the deep past. Two haplotypes (single sample) per region were used, thus four haplotypes (4 x 7 = 28 chromosomal haplotypes) in total, in cross-population analyses. Instead, multiple sample pairs between regions were compared to confirm robustness of results. This approach, coupled with the high-resolution timing of demographic events aligned with marine isotope stages (Fig. 3, fig. S10-13, Text S1), negated the need for conducting additional bootstrap analyses for individual pairs.

To scale historic events for the woodland strawberry, the experimentally tested mutation rate of *A. thaliana* of 7.1*10^-9^ per bp per generation (Ossowski et al. 2010) and a two-year generation time that has been used for studies on other perennial herbs (Savolainen and Kuittinen 2011, Koch et al. 2006) were used. Using the MSMC-IM the time-dependent (generations ago) symmetric migration rate (m(t)) and migration rate refined coalescence rates (effective population size, N_E_) were inferred for each population pair (im_N1 and im_N2). im_N1 was selected to represent historical N_E_ because the analyses with some sample pairs finished successfully without clear errors only when a specific sample was assigned as im_N1. This was especially the case when one of the samples had a small current N_E_. All MSMC-IM output files (designated as *.estimate-files) underwent a thorough review (Table S3). Strong glacial periods, such as MIS 6 (or PGP), often led to failures of runs (8% of all runs) due to extreme bottlenecks in either one or both populations. Those runs were not accepted for estimating initial divergence (M<0.999, m<0.99), but were accepted for estimating split times (estimated median split time of a population pair, M(t) < 0.5) that occurred during the Holocene. If a run failed during the Holocene, split times were not accepted; however, those runs were accepted to estimate initial divergence, which usually initiated much earlier in history. If a failure occurred more than once, leading to fluctuations of N1 and N2 parameters between extreme values, runs were discarded from results. Such runs often occurred when the current effective population sizes (N_E_) were low in both populations (indicated by low im_N1 and im_N2 values, as observed in the Alta-Kåfjord populations). However, for assessing current N_E_, both im_N1 and im_N2 parameters were accepted, given that earlier time points do not have an impact (the principle of Markovian property) on the most recent coalescence rates (Table S2). As mentioned earlier, hybrid samples with strongly negative F_IS_ values (ES21, ES3, NOR28, NOR30) were excluded from regional pools. Additionally, two Iberian samples (ES13 and ES18) that showed suspiciously steep recent growth (< 650 generations ago) despite still being highly inbred were excluded.

### Heterozygosity

Nucleotide heterozygosity for each sample was estimated with ANGSD-software (Korneliussen et al. 2014) directly from prefiltered bam-files using the folded frequency spectrum. The following filters were used: uniqueOnly 1, minQ20, minmapQ 30 -C50. Regional nucleotide diversities were calculated from imputed and phased polymorphic sites using vcftools (Danecek et al. 2011). Intergenic sites were used to compare nucleotide diversities between regions.

### Data, Materials, and Software Availability

Genome sequence data have been deposited to NCBI (PRJNA1018297; will be made publicly available after the acceptance of the manuscript). The code used for data-analysis is available at Github (https://github.com/tuomas64/strawberry).

## Acknowledgements

The project was funded by European Commission (BiodivERsA project BiodivClim-177) to T.H., the Academy of Finland (grant nr 317306 and 344726 to T.H. and grant nr 315353 to T.T.), and European Research Council (ERC-2014-StG 638134) and the Spanish Ministries of Science, Innovation and Universities (PID2021-123677OB-I00) to D.P. The study included material from the “Professor Staudt Collection” maintained in Dresden, Germany at the Hansabred GmbH & Co. KG and from the IFAPA *Fragaria* collection funded by IFAPA Project PR.CRF.CRF202200.002 of the European Agricultural Fund for Rural Development, within the Rural Development Program of Andalusia 2014-2022. We thank Eija Takala and Marjo Kilpinen for laboratory assistance and Katriina Palm for maintaining the plant materials.

