## Supplemental text, figures and table for "Late Quaternary climatic impact on the woodland strawberry genome: a perennial herb’s tale"

**This PDF file includes:**

Supporting text S1 and S2
Figures S1 to S21
Table S1

### Supporting Information Text

#### Text.S1. High resolution MSMC-IM analysis in samples with low $F_{IS}$ and high $N_E$

For historical $N_E$ and migration rates, we focused on common clear patterns (Fig. S17), that aligned with the chronological timing of serial MIS (Lisiecki and Raymo 2005). High resolution MSMC-IM runs often occurred when one sample was drawn from an eastern core population (Croatia, Lithuania, or Romania) or another population with high $N_E$ because they best met the expected conditions for long-term sexual reproduction. Especially in the peripheral regions, samples with the largest $N_E$ and the lowest $F_{IS}$ gave the most accurate temporal resolution, probably because large segments of their genomes were more recently diverged from their source populations with high $N_E$ (Fig. S8, S9), while the analysis of the smallest populations often failed or resulted in low resolution histories. Demographic history patterns were divided in different categories according to following criteria (Fig. S17, Table S3): CORE-1=Stable migration rate from MIS 11 until LGM, strongly increased Ne during MIS 11, strong bottleneck during MIS 12, slightly reduced Ne during PGP and possibly during MIS 16; CORE-2=Stable migration rate from MIS 11 until LGM, slightly reduced Ne during PGP; PERIPHERAL-1=One strong bottleneck in addition to the LGM bottleneck during either MIS 6 or MIS 8, fluctuating migration rate at least during two G-IG cycles between PGP and MIS 10; PERIPHERAL-2=Stable $N_E$, migration completely terminated at least once between MIS 10 and LGM; PERIPHERAL-3=One strong bottleneck and termination of migration rate at least once between MIS 10 and LGM; PERIPHERAL-4=Strong bottleneck between MIS 10 and PGP, no termination of migration between MIS 10 and LGM. In our analyses, we focused on CORE-1 and PERIPHERAL-1 patterns. Results of all MSMC-IM runs are available in Table S3.

#### Text S2. Reliability of migration rates

In MSMC-IM runs, sample pairs with a cumulative migration probability (M) lower than 0.999 (Fig. S14, S17A, S21), a threshold deemed acceptable for detecting migration rates between two ancestral lineages according to Wang et al. (2020), exhibited migration patterns identical to those above the threshold (M>0.999, Fig. S10-13, S15-16). Specifically, both showed a complete cessation of migration during the MIS 12 (CORE 1) and MIS 10, MIS 8, MIS 6 (PERIPHERAL 1) glacial periods and more recently during the LGM. There can be several reasons for that. In the original paper in humans (Wang et al. 2020), the maximum age of initial divergence (M<0.999) was about 60 000 generations, after which two lineages were fully randomized. In this study, we found consistent patterns for up to 165,000-185,000 generations (330-370 ka, corresponding to MIS 10) in the peripheral pattern, indicating higher temporal resolution limit in woodland strawberry compared to human. Aligning with high temporal resolution of the peripheral pattern, several population pairs began to diverge (M<0.999) during MIS 10 (Fig. S7). For example, four independent sample pairs between Iberia and Italy (different refugia) started to diverge (M<0.999) during that time. In the core pattern, temporal resolution extended up to 300,000 generations (600 ka, corresponding to MIS 16). However, distant initial divergence events (M<0.999) occurred less frequently, with only a few population pairs between Italy/Croatia and Romania or Lithuania and Romania, initiating divergence during or before MIS 16 or MIS 12. Consequently, there is greater uncertainty regarding the migration rates in the core pattern.

Both genomic and biological reasons can contribute to this difference between strawberries and humans. In small genomes like in woodland strawberry (~220Mb), where genic regions comprise nearly half of the genome, constructing accurate haplotype data may be relatively straightforward. Moreover, woodland strawberries, particularly in peripheral areas, likely reproduced by selfing during glacial maxima, as our results suggest in current peripheral populations. This led to a complete cessation of migration between separate refugia. Subsequently, these selfed lineages hybridized with other populations during interglacial range expansions, when hybrids possibly gained a fitness advantage. This is possible reason why we observed fluctuating migration rate pattern so frequently in peripheral samples (about 30% of all successful runs). It is possible that the varying time-dependent modes of reproduction and cycling between ancestral homozygosity and heterozygosity, are more readily detectable in strawberry genomes than in human genomes, where self-fertilization does not occur. Extensive hybridizations, particularly during the Holocene, are evident in strawberry genomes. Sample pairs showing less hybridization during the Holocene, such as between Iberia and Lithuania/Romania, which split during the MIS2, began diverging

(M<0.999) consistently in a deeper past (Fig. S7). This supports our hypothesis that extensive hybridizations during the Holocene have often increased the cumulative migration probability threshold above 0.999, despite the possibility that old parts of ancestral lineages can still be distinguishable. These factors suggest that studies on strawberry genomes might reveal deeper initial divergences (M>0.999) compared to those observed in human populations.

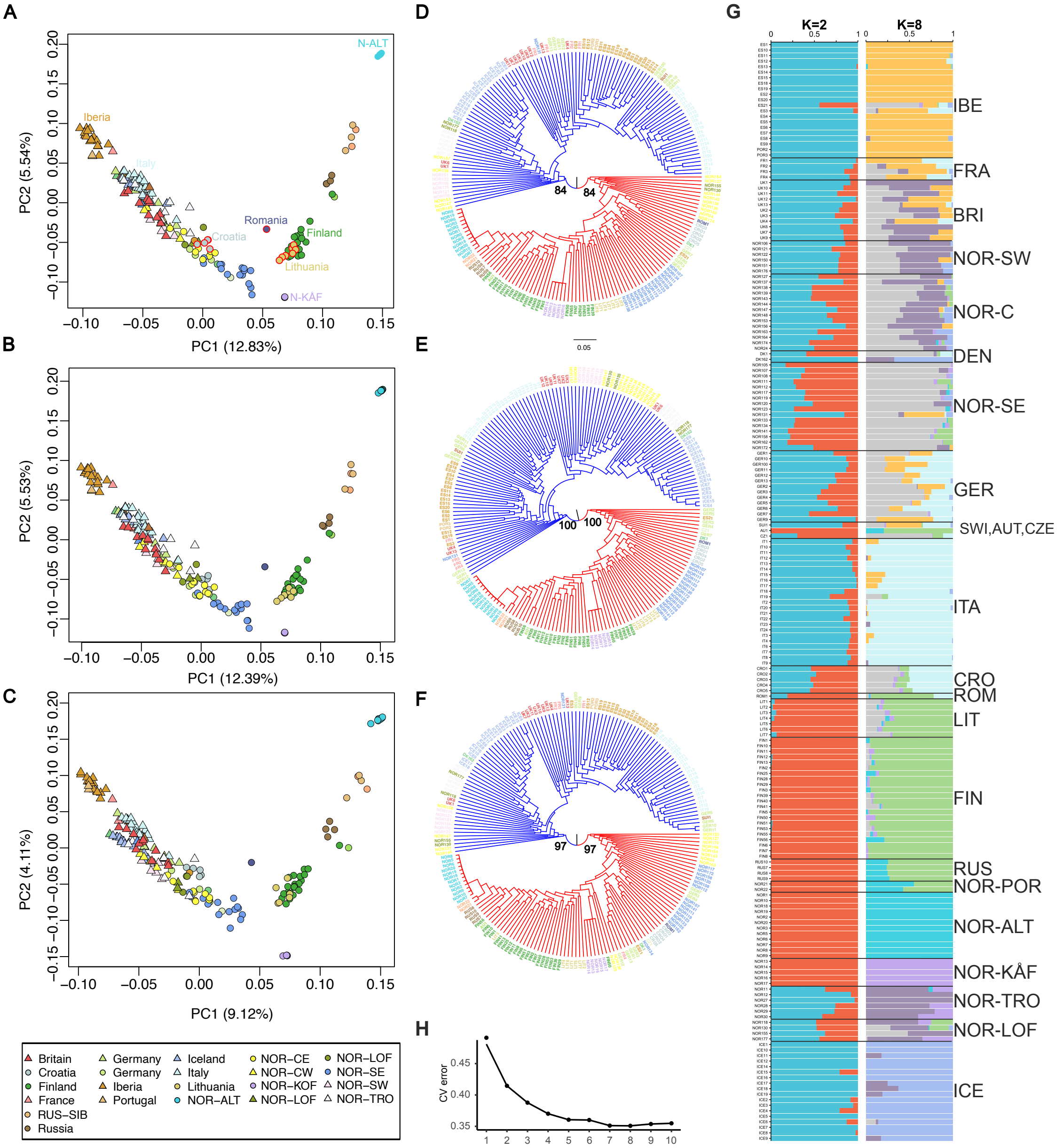


Fig. S1. Population structure using different genomic datasets and analyses. A-C) PCA using (A) 4-fold degenerate sites, (B) nonsynonymous sites and (C) LD-pruned set of SNPs across the whole genome. Bootstrap values for western and eastern branches are shown. D-F) Hierarchical clustering (HC) based on the same datasets. G) Admixture plot with K=2 and K=8 using 4-fold degenerate sites. H) Cross-validation error for different values of K. Abbreviations: IBE=Iberia, FRA=France, BRI=Britain, NOR-SW=Norway-southwestern, NOR-C=Norway-central,

DEN=Denmark, NOR-SE=Norway-southeastern, GER=Germany, SWI=Switzerland, AUT=Austria, CZE=Czechia, ITA=Italy, CRO=Croatia, ROM=Romania, LIT=Lithuania, FIN=Finland, RUS-NW=Russia-northwestern, NOR-POR=Norway-Porsanger, NOR-ALT=Norway-Alta, NOR-KÅF=Norway-Kåfjord, NOR-TRO=Norway-Tromso, NOR-LOF=Norway-Lofoten, ICE=Iceland, SWI=Switzerland.

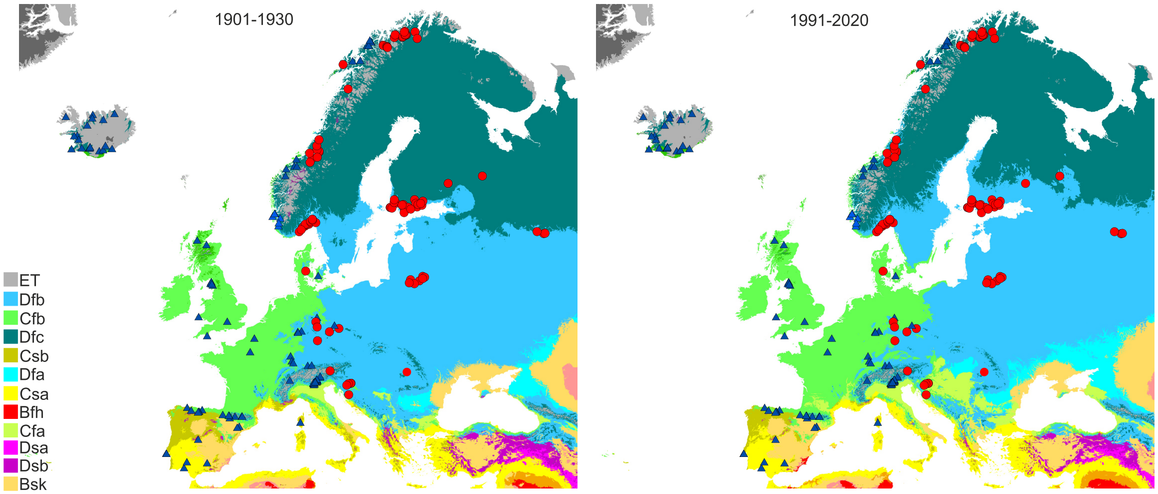


Fig. S2. Geographical division of the western and eastern genetic clusters of woodland strawberry (based on maximum likelihood clustering at 4-fold degenerate sites) follows the border separating oceanic (Cfb) and continental (Dfb) or sub-arctic (Dfc) climatic zones across Europe (Beck et al. 2023). Modeling the distributions of climatic zones for two different time-intervals: A) (1901-1930) and B) (1991-2020). Other climatic zones: ET=Tundra, Csb=Warm-summer Mediterranean climate, Dfa=Hot-summer humid continental climate, Csa=Hot-summer Mediterranean climate, Bfh=Hot desert climate, Cfa=Humid subtropical climate, Dsa = Mediterranean-influenced hot-summer humid continental climate, Dsb = Mediterranean-influenced warm-summer humid continental climate, BSk = Cold semi-arid climate.

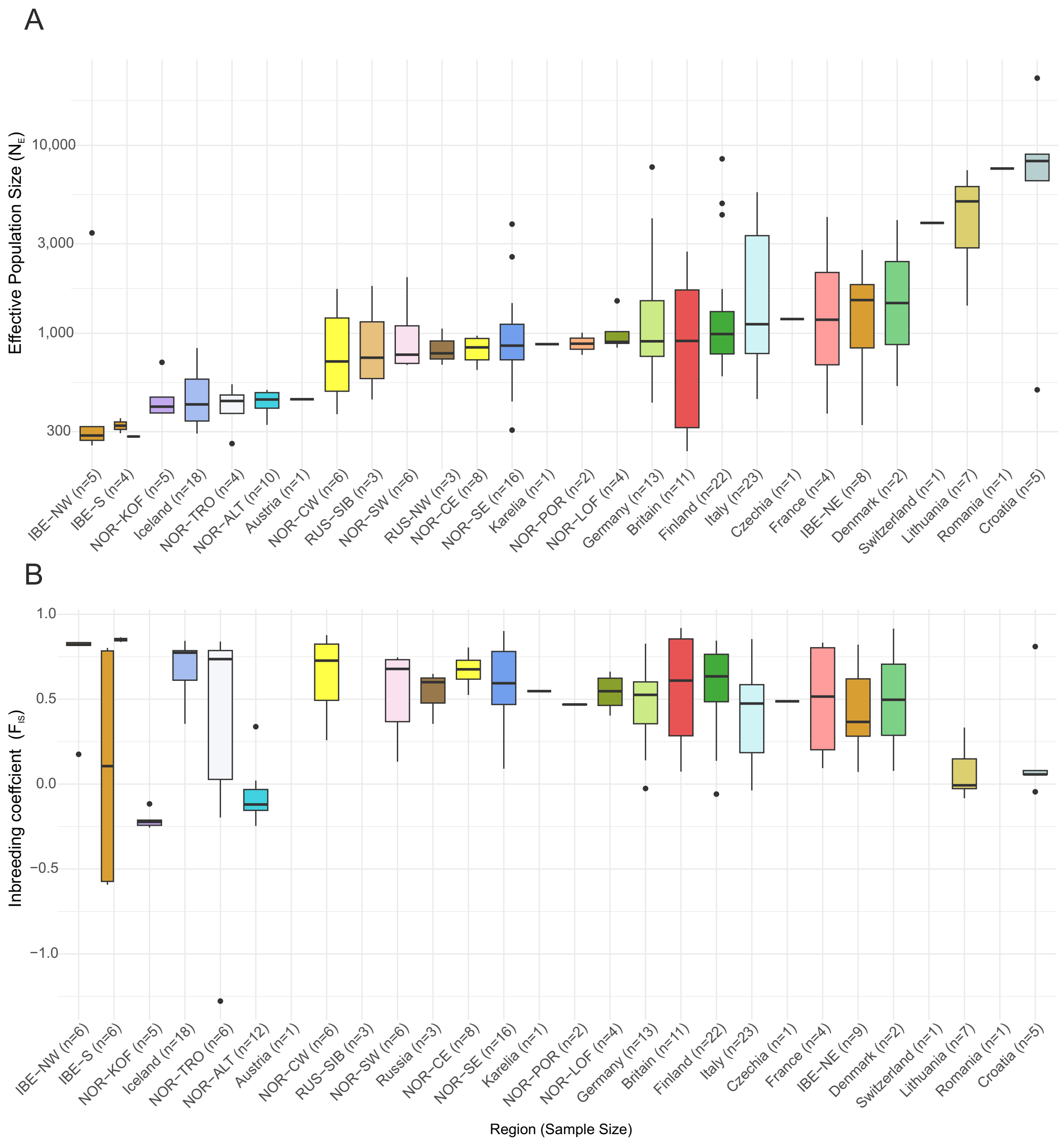


Fig.S3. Effective population size (A) and regional inbreeding coefficients (B) ordered by increasing $N_E$. Numbers at the bottom of figures show the number of samples used in corresponding figures. Note that $F_{IS}$ cannot be calculated if sample size is n=1. One sample from Karelia (RUS10) and Czechia (CZ1) were combined for their $F_{IS}$ estimation with three samples from northwest Russia (Russia) and German samples, respectively. Abbreviations: IBE-S=southern Iberia, IBE-NW=northwestern Iberia, NOR-TRO=Norway-Tromso, NOR-KÅF=Norway-Kåfjord, NOR-POR=Norway-Porsanger, NOR-ALT=Norway-Alta, NOR-CW=Norway-central-western, NOR-SE=Norway-southeastern, NOR-LOF=Norway-Lofoten, NOR-SW=Norway-southwestern, NOR-CW= Norway-central-eastern, IBE-NE=, northeastern Iberia.

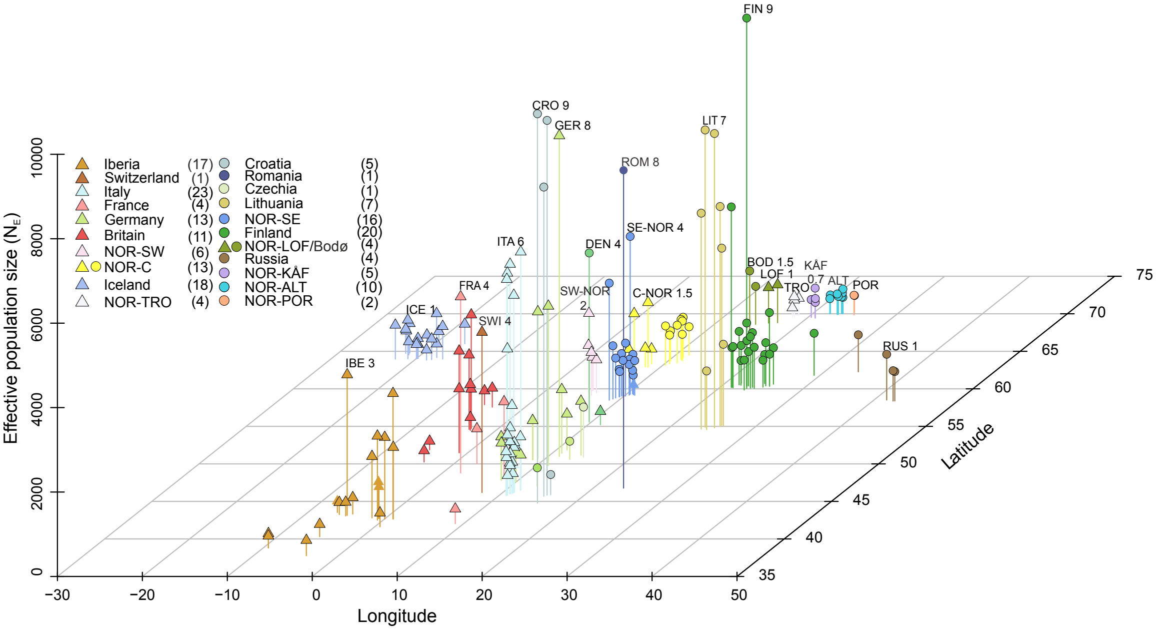


Figure S4. Populations at the edges of the range have the lowest effective population sizes ($N_E$). $N_E$-values plotted with geographic coordinates of each sample. Population with the highest $N_E$ in each region is highlighted.

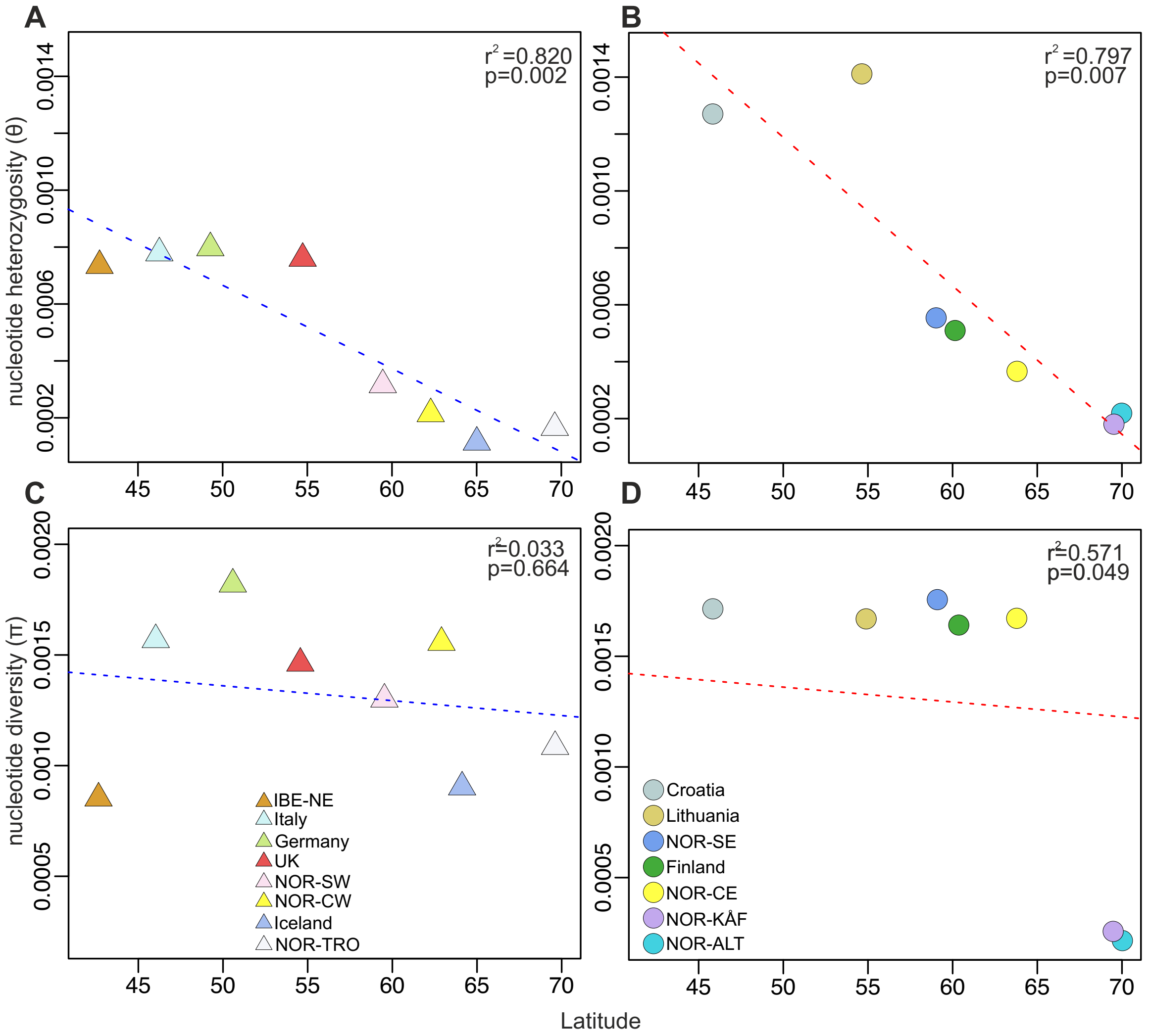


Figure S5. Latitudinal correlation of genetic diversity. Median nucleotide heterozygosity (theta) per sample in A) western and B) eastern regions. Intergenic nucleotide diversity (π) per region in C) western and D) eastern Europe. The coefficient of determination ($r^2$) is derived from the Pearson correlation coefficient (r), with statistical significance indicated by the *p*-value.

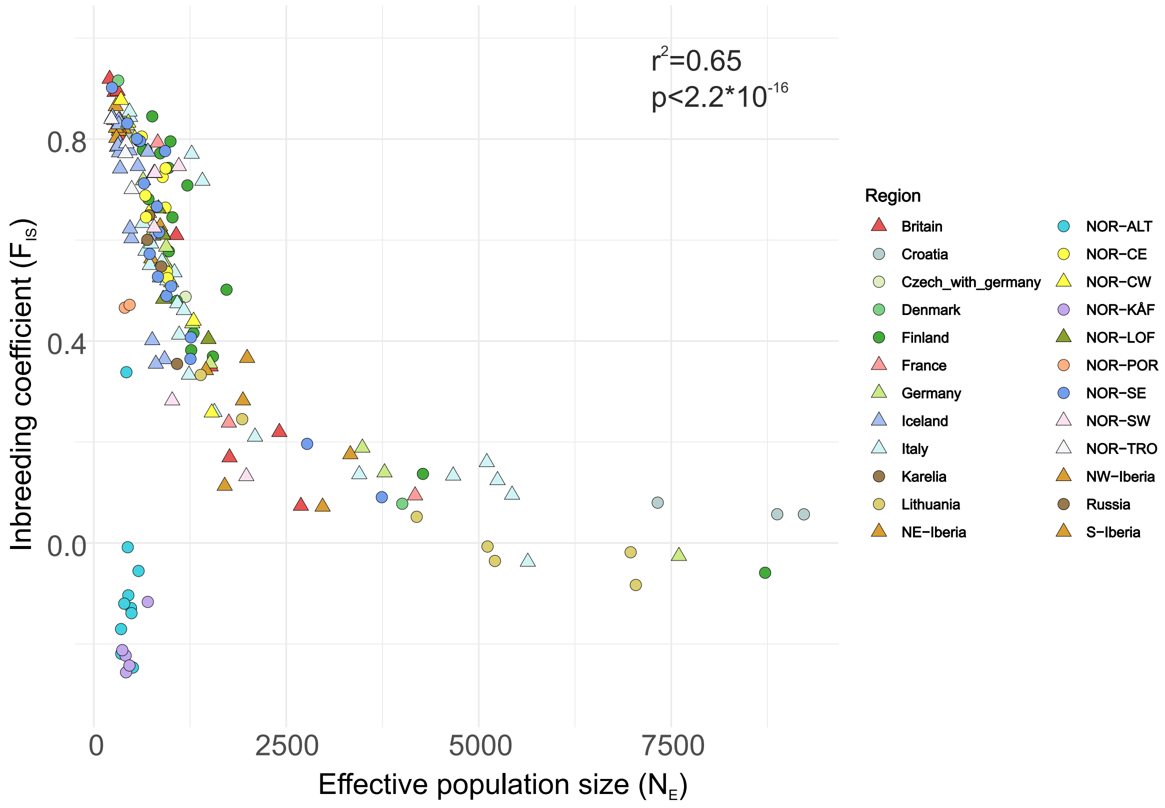


Figure S6. Correlation of regional inbreeding coefficients and present effective population sizes. The coefficient of determination ($r^2$), and a significance of correlation were calculated excluding Alta (NOR-ALT) and Kåfjord (NOR-KÅF) that clearly differed from the general pattern.

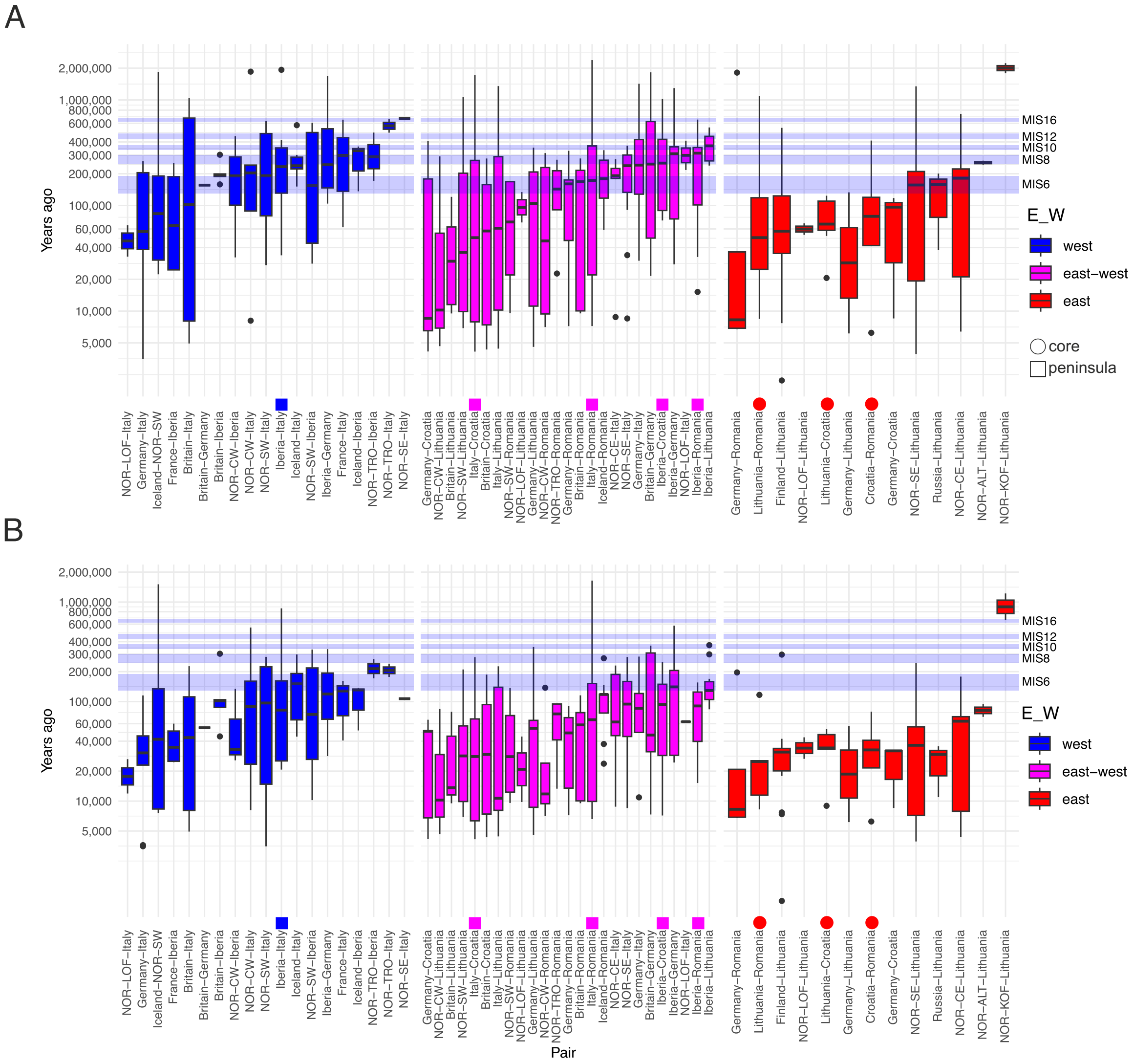


Fig. S7. Initial divergence (ya) between regional pairs of samples with A) (M<0.999) and B) (M<0.99). Circles after the sample pair names highlight initial divergence between core populations and rectangles between southern European Peninsulas. Glacial periods (MIS6-MIS16) shown by blue shading.

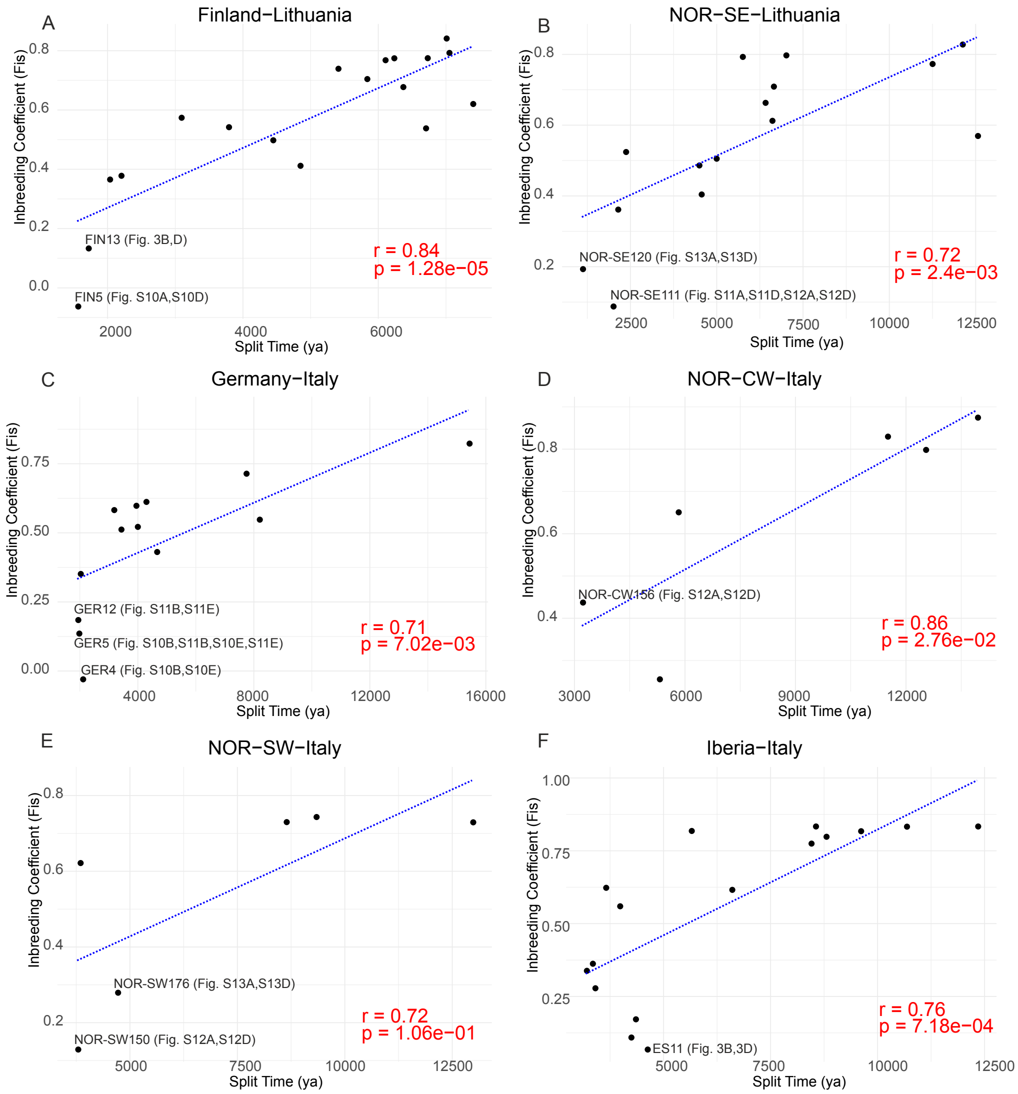


Figure S8. Peripheral populations with the lowest inbreeding coefficients (hybrids) split the most recently from their source populations. Correlation of split times and inbreeding coefficients ($F_{IS}$). $F_{IS}$ of the first population mentioned in the figure title was plotted against the split time of the population pair. r=correlation coefficient, p=p-value based on cor.test function in R. Samples with high quality histories in specific figures were highlighted.

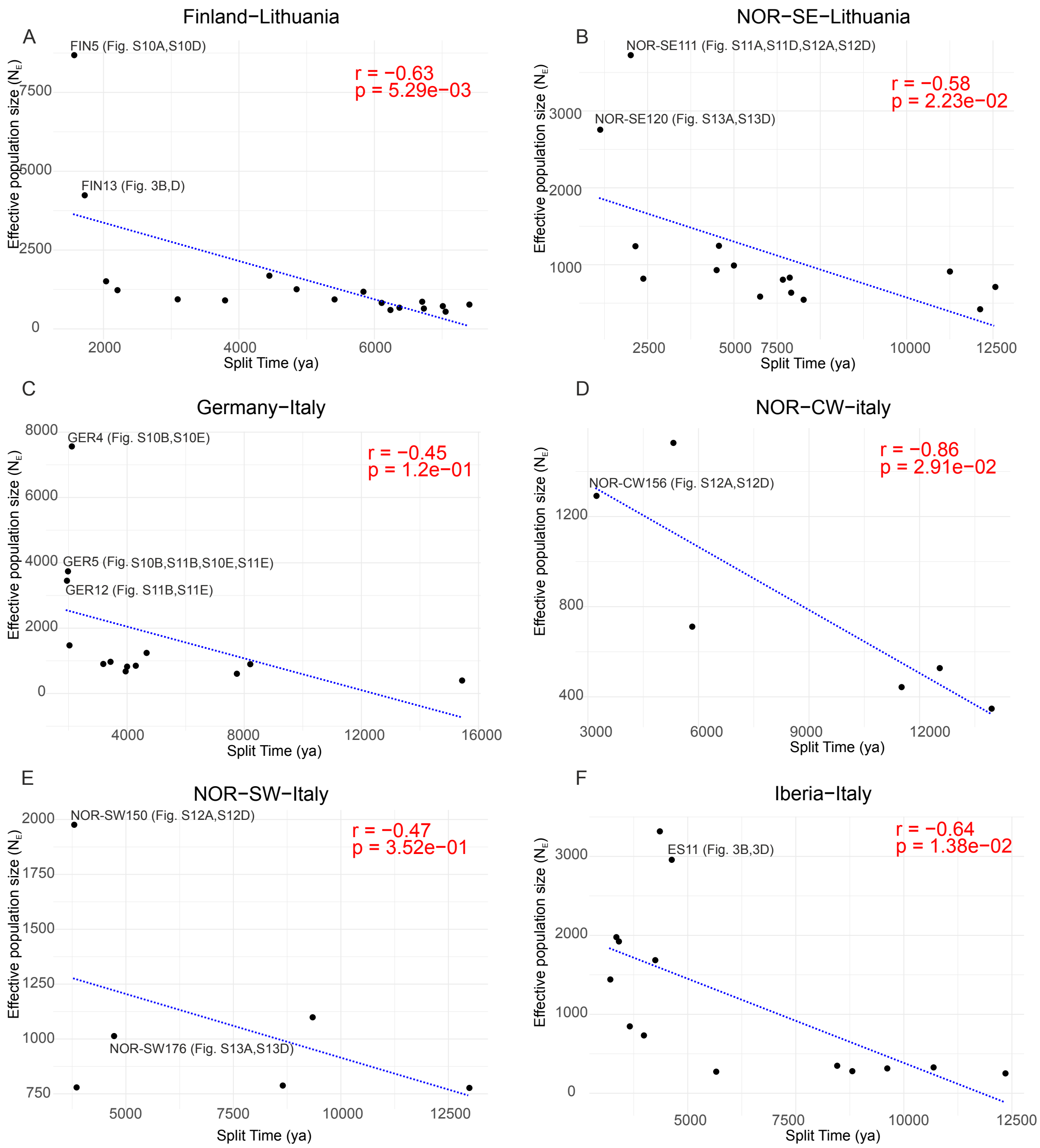


Figure S9. The largest peripheral populations split the most recently from their source populations. Correlation of split times and effective population sizes ($N_E$). $N_E$ of the first population mentioned in the figure title was plotted against the split time of the population pair. r=correlation coefficient, p=p-value based on cor.test function in R. Samples with high quality histories in specific figures were highlighted.

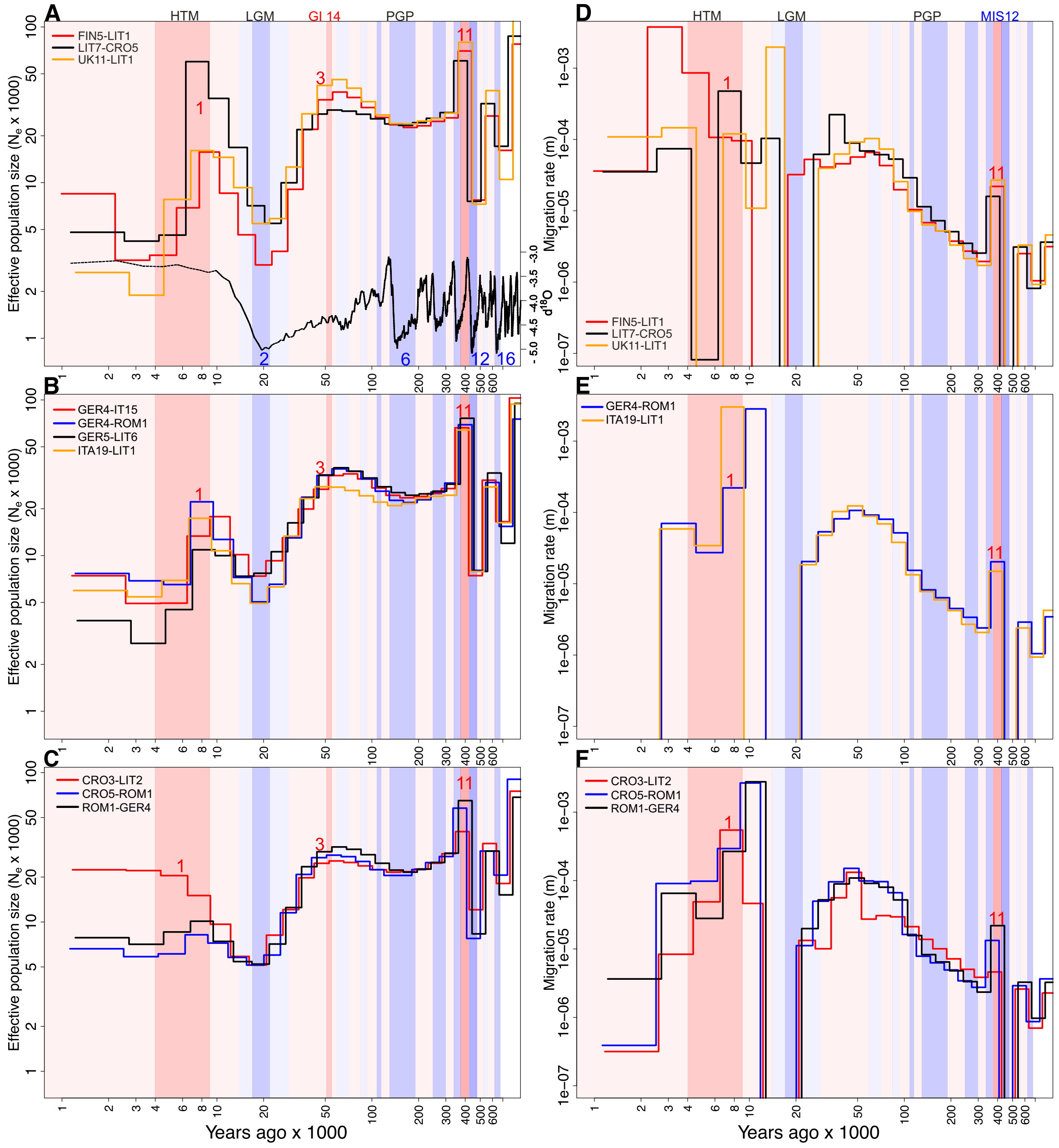


Fig. S10. Climatic histories in the core pattern (CORE 1) with the history extending until MIS16 with acceptable resolution. A-C) Effective population sizes ($N_E$) and D-F) migration rates of the core patterns of history. HTM=Holocene Thermal Maximum (9,000-4,000 ya), LGM=Last Glacial Maximum (22,000-17,000 ya) GI14=Greenland Interstadial 14 (55,000-51,000 ya) PGP=Penultimate Glacial Period (190,000-130,000 ya). Curve at the bottom of A represents inverse benthic δ18O records from Lisiecki and Raymo (2005) and is used as a proxy for historical temperature.

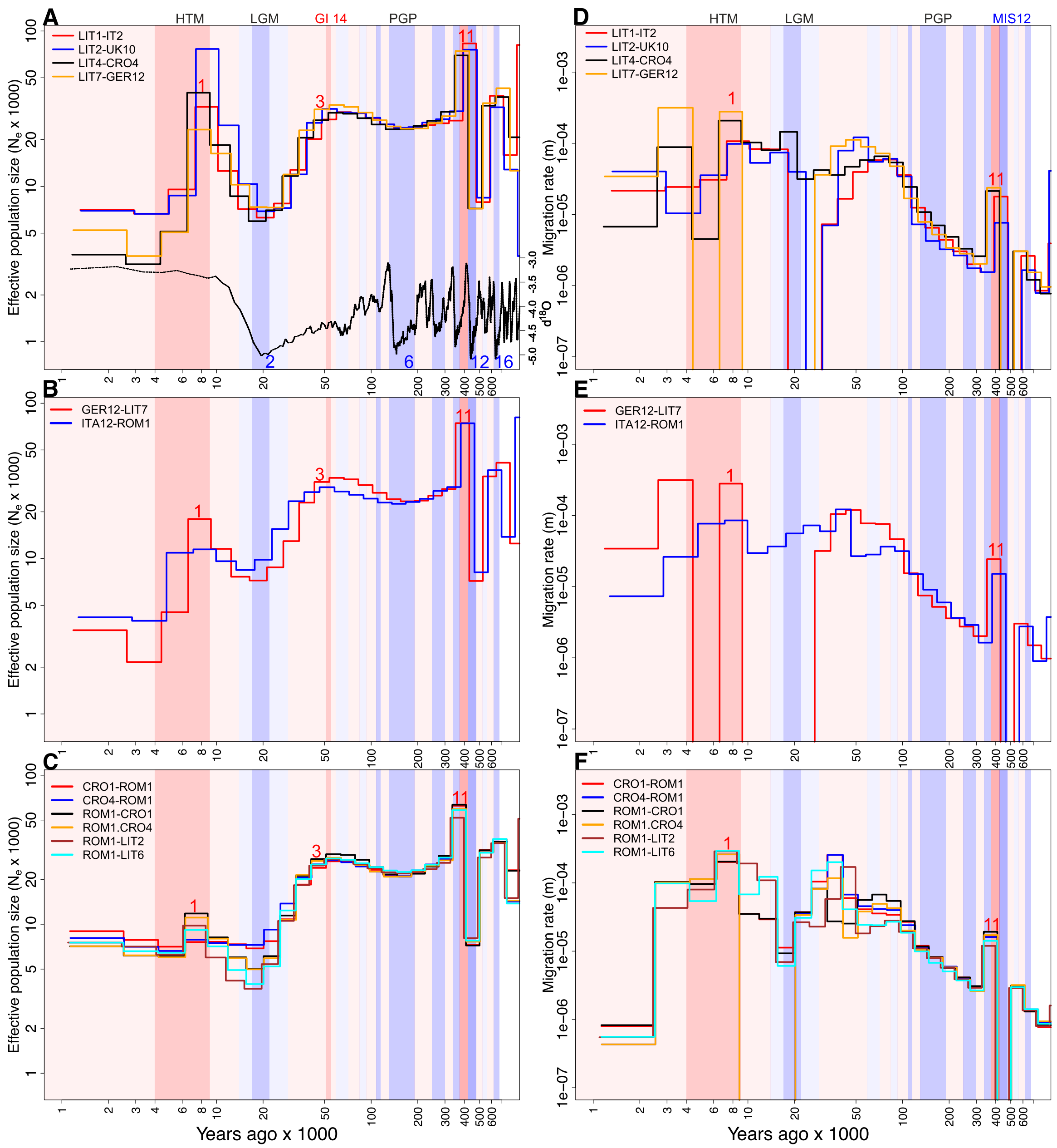


Figure S11. Climatic histories in the core pattern (CORE 1) with the history extending until MIS12 with acceptable resolution. A-C) Effective population sizes ($N_E$) and D-F) migration rates of the core patterns of history. HTM=Holocene Thermal Maximum (9,000-4,000 ya), LGM=Last Glacial Maximum (22,000-17,000 ya), GI14=Greenland Interstadial 14 (55,000-51,000 ya), PGP=Penultimate Glacial Period (190,000-130,000 ya). Curve at the bottom of A represents inverse benthic δ18O records from Lisiecki and Raymo (2005) and is used as a proxy for historical temperature.

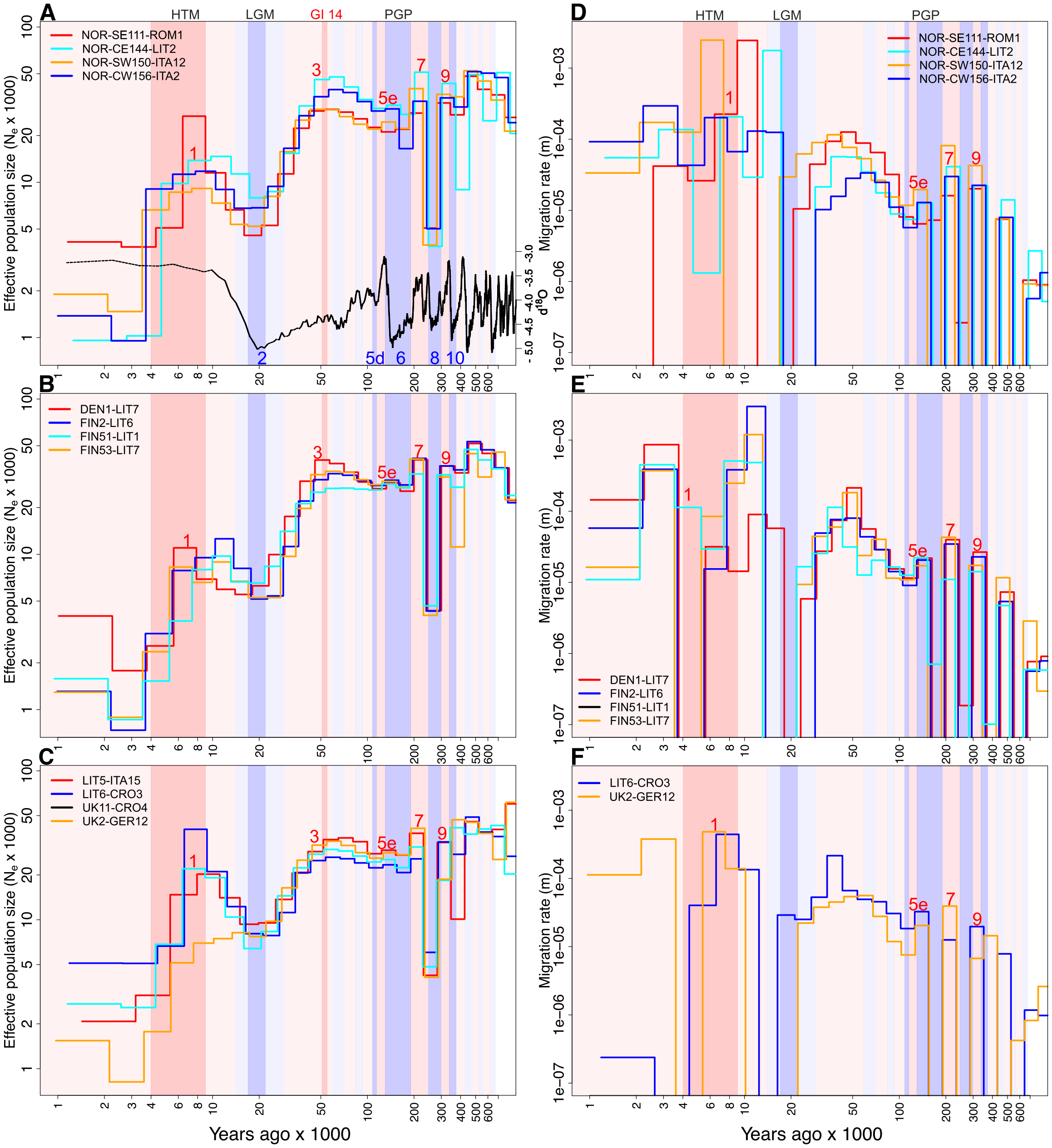


Figure S12. Climatic histories in peripheral pattern (PERIPHERAL 1), showing strong bottlenecks during MIS8, with the history extending until MIS10 with acceptable resolution. A-C) Effective population sizes ($N_E$) and D-F) migration rates of the peripheral pattern of history. HTM=Holocene Thermal Maximum (9,000-4,000 ya), LGM=Last Glacial Maximum (22,000-17,000 ya), GI14=Greenland Interstadial 14 (55,000-51,000 ya), PGP=Penultimate Glacial Period (190,000-130,000 ya). Curve at the bottom of A represents inverse benthic δ18O records from Lisiecki and Raymo (2005) and is used as a proxy for historical temperature.

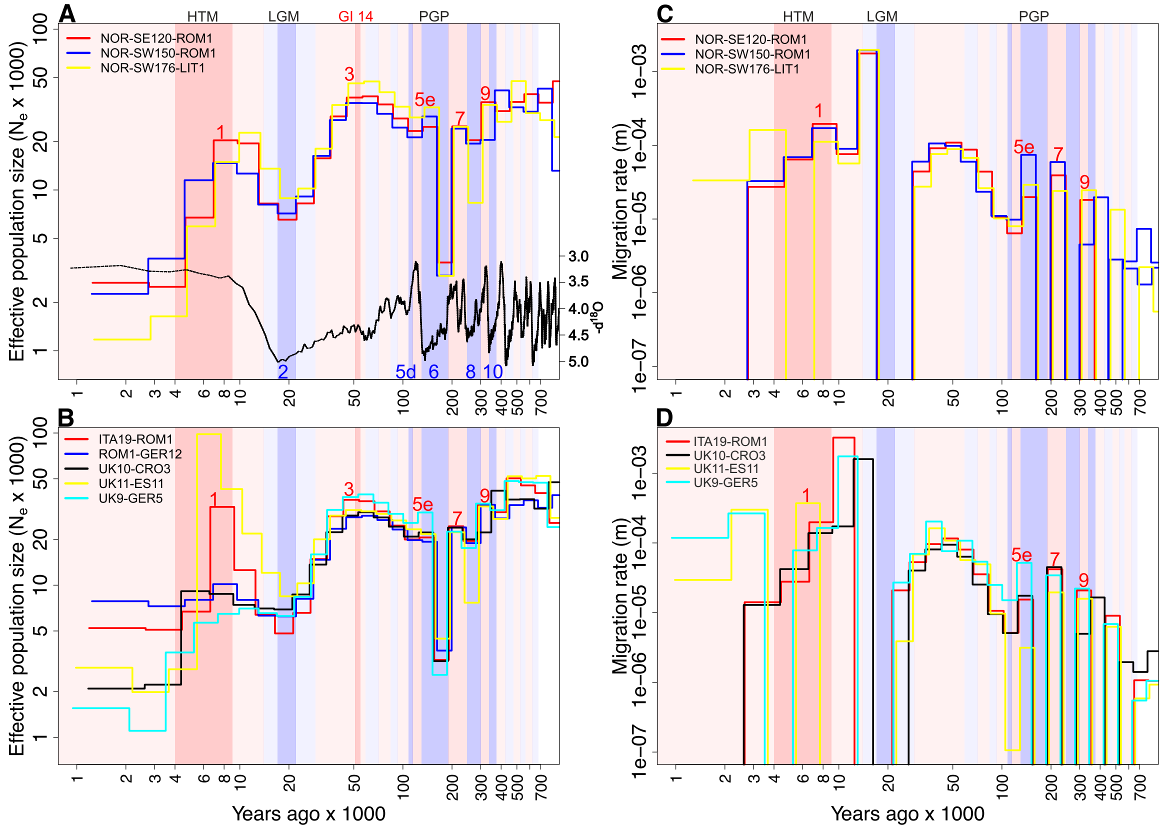

Figure S13. Climatic histories in peripheral pattern (PERIPHERAL 1), showing strong bottlenecks during MIS6, history extending until MIS10 with acceptable resolution. A-C) Effective population sizes ($N_E$) and D-F) migration rate of the peripheral pattern of history. HTM=Holocene Thermal Maximum (9,000-4,000 ya), LGM=Last Glacial Maximum (22,000-17,000 ya), GI14=Greenland Interstadial 14 (55,000-51,000 ya), PGP=Penultimate Glacial Period (190,000-130,000 ya). Curve at the bottom of A represents inverse benthic δ18O records from Lisiecki and Raymo (2005) and is used as a proxy for historical temperature.

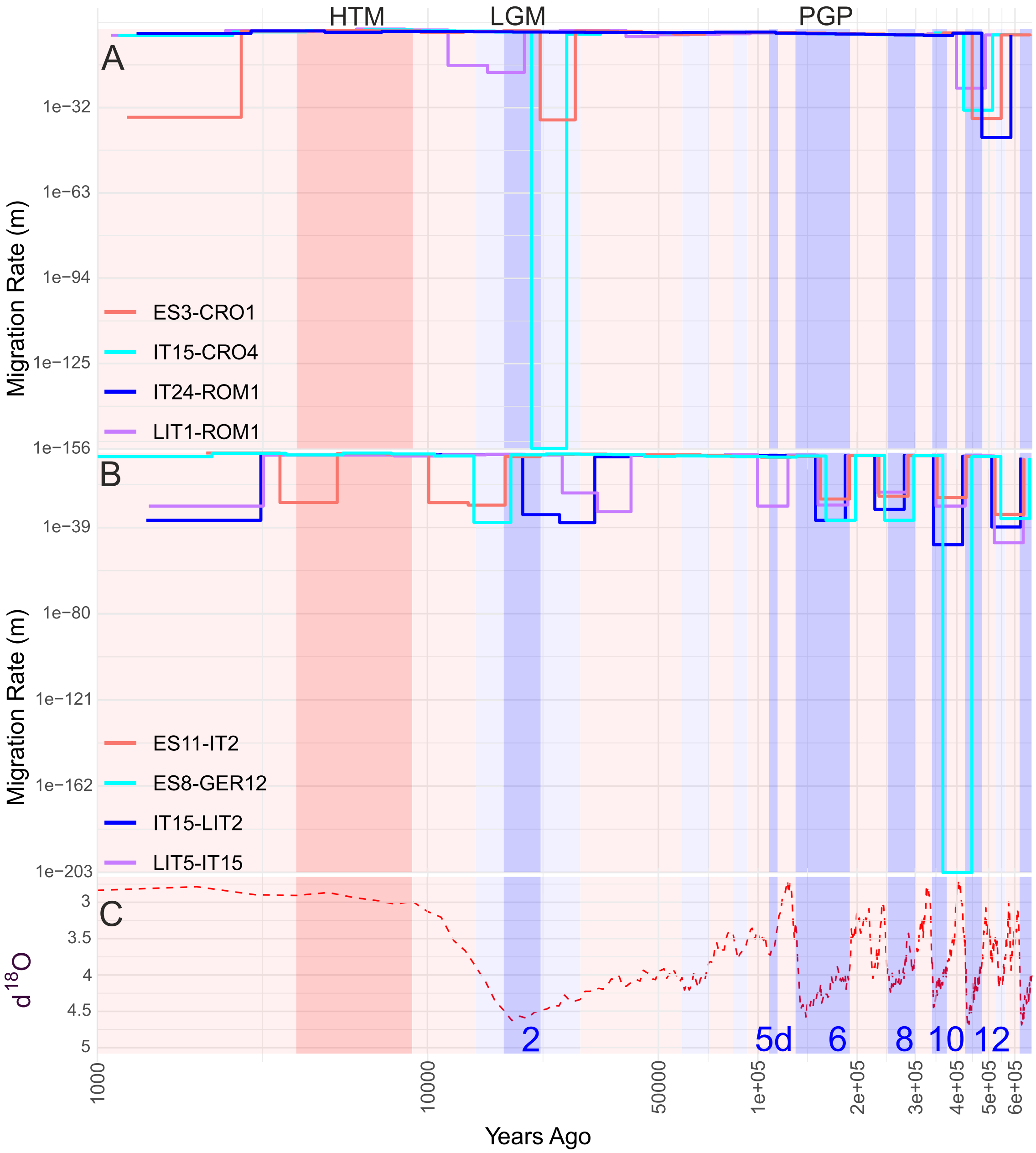


Fig. S14. Symmetric migration rate through time. A) Core (CORE 1) and B) peripheral patterns (PERIPHERAL 1) of history. The five most recent major glacial periods are shown by numbers for specific Marine Isotope Stages (2=MIS2, 6=MIS6, 8=MIS8, 10=MIS10, 12=MIS12). M<0.999 from MIS16 (A) and from MIS10 (B) until present time. C) The δ18O isotope data from Lisiecki and Raymo (2005).

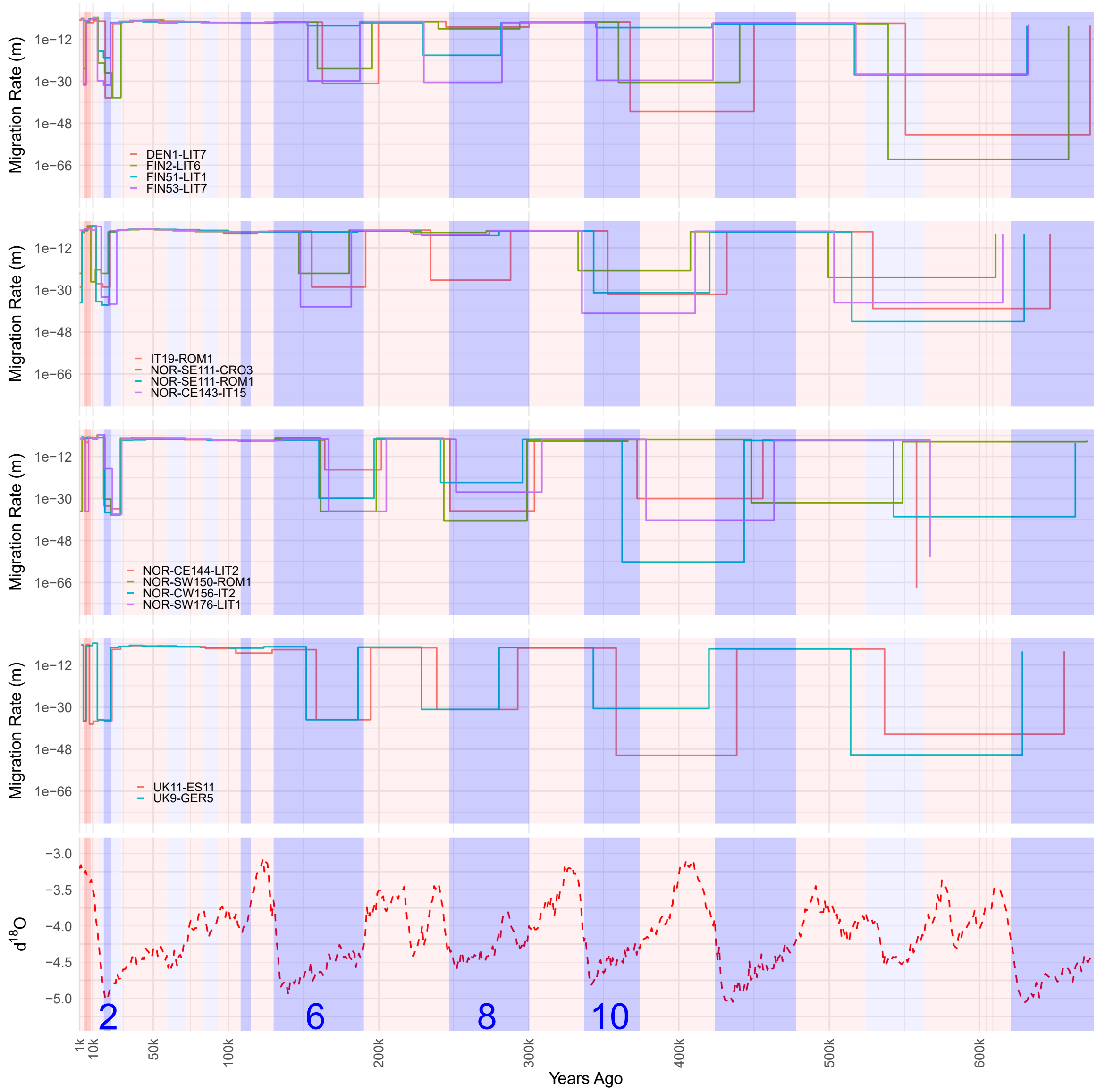


Fig. S15. Symmetric migration rate through time. Peripheral patterns of history are shown using linear scale. The four most recent major glacial periods shown by numbers for specific Marine Isotope Stages (2=MIS2, 6=MIS6, 8=MIS8, 10=MIS10). The lowest panel shows the δ18O isotope data from Lisiecki and Raymo (2005).

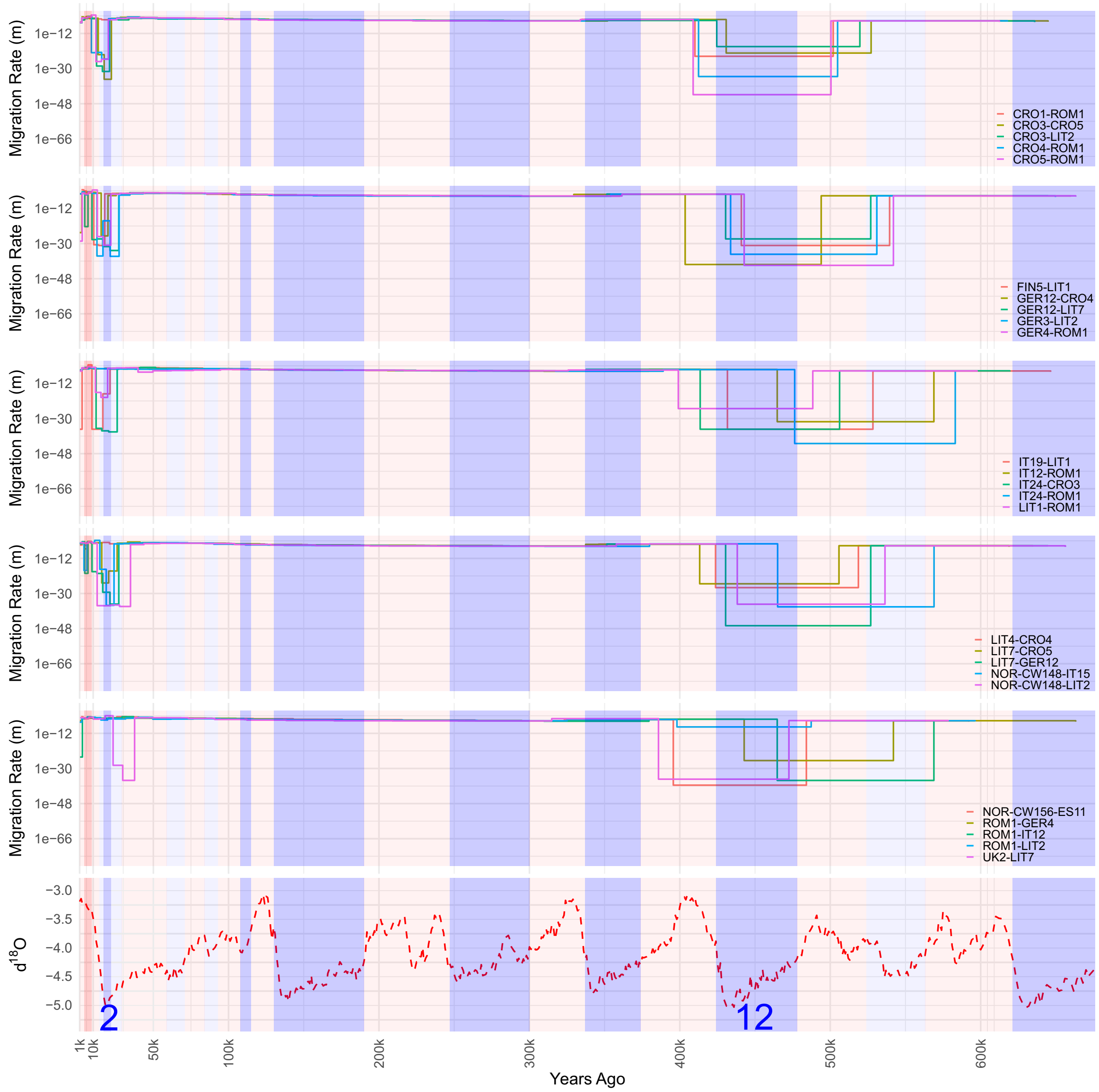


Fig S16. Symmetric migration rate through time. Core patterns of history using linear scale of time. Glacial periods shown by numbers for specific Marine Isotope Stages (2=MIS2,12=MIS12). The lowest panel shows the δ18O isotope data from Lisiecki and Raymo (2005).

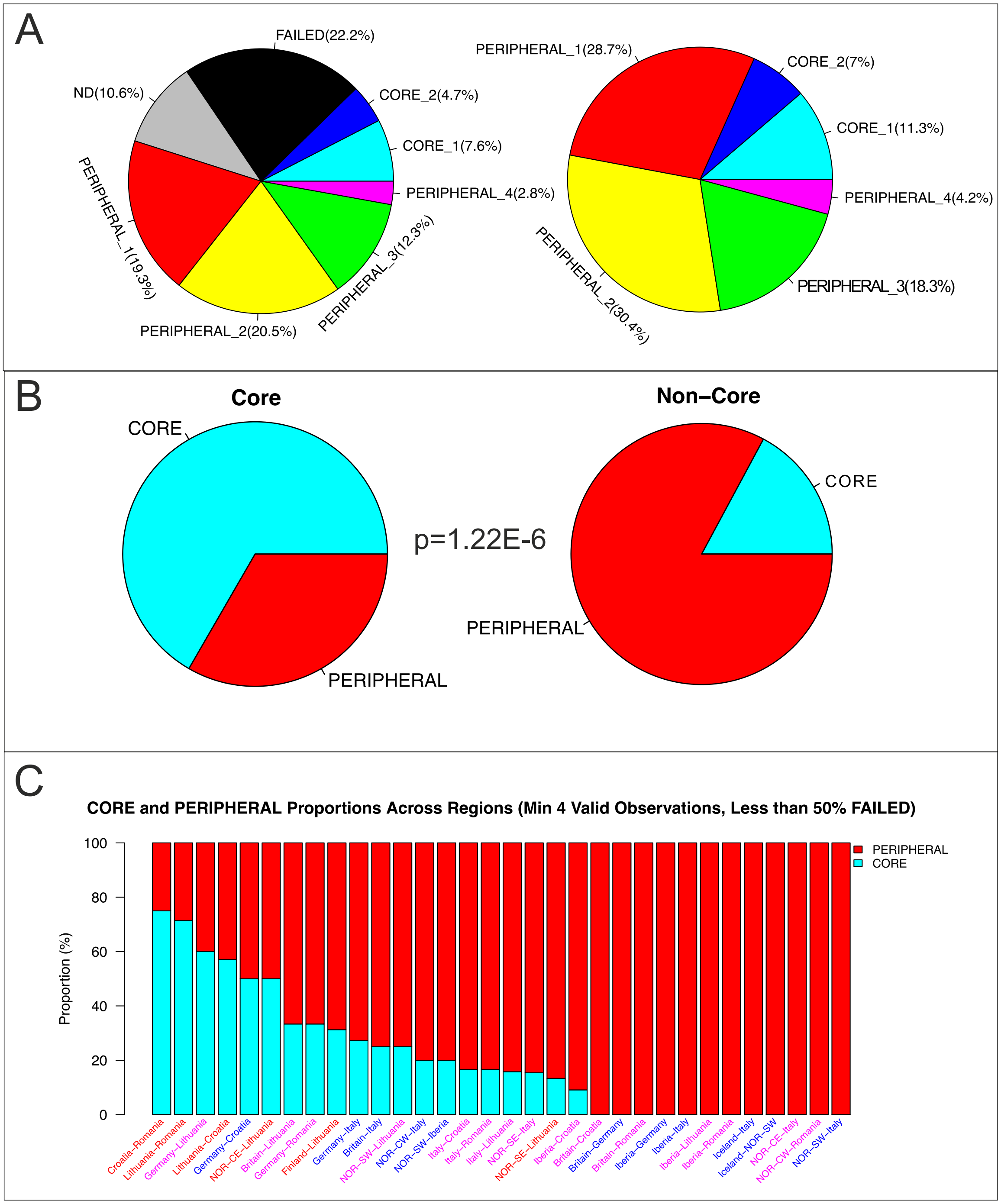


Fig. S17. A) Proportion of different demographic history patterns. The pie in left includes all sample comparisons, while in right, FAILED and ND patterns are excluded. B) Proportions of core (CORE1+CORE2) and peripheral (PERIPHRAL1-4) patterns in the comparisons of core samples (Croatia-Romania, Lithuania-Croatia and Lithuania-Romania haplotype combinations) and in other comparisons (Non-Core). The Fischer exact test was used to calculate *p*-value for the difference. C) Proportion of core and peripheral patterns in different sample pairs.

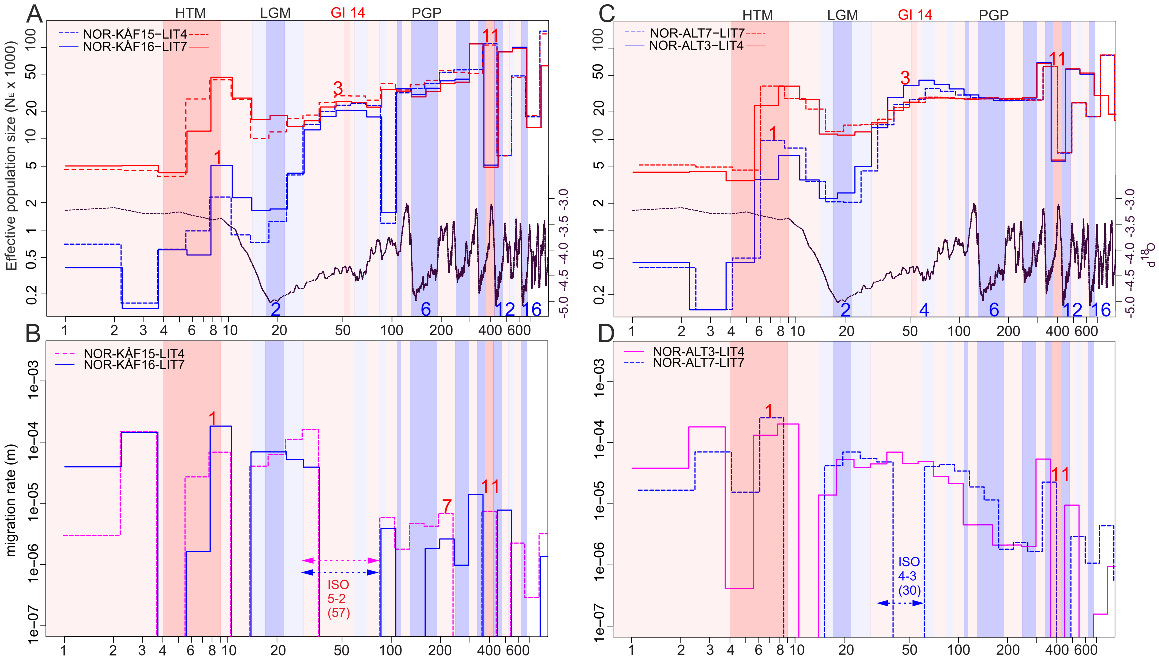


Figure S18. Demographic histories of northern Norwegian Kåfjord and Alta populations. A) Effective population sizes of Kåfjord (blue) and Lithuania (red). B) Migration rate between Kåfjord and Lithuania. C) Effective population size of Alta (blue) and Lithuania (red). D) Migration rate between Alta and Lithuania. Glacial and interglacial periods are shown by blue and red background shading and numbers. ISO=isolation event. Numbers in parentheses describe lengths of isolation events in thousands years. HTM=Holocene Thermal Maximum (9,000-4,000 ya), LGM=Last Glacial Maximum (22,000-17,000 ya), GI14=Greenland Interstadial 14 (55,000-51,000 ya), PGP=Penultimate Glacial Period (190,000-130,000 ya). Curve at the bottom of A and B represents inverse benthic δ18O records from Lisiecki and Raymo (2005) and is used as a proxy for historical temperature.

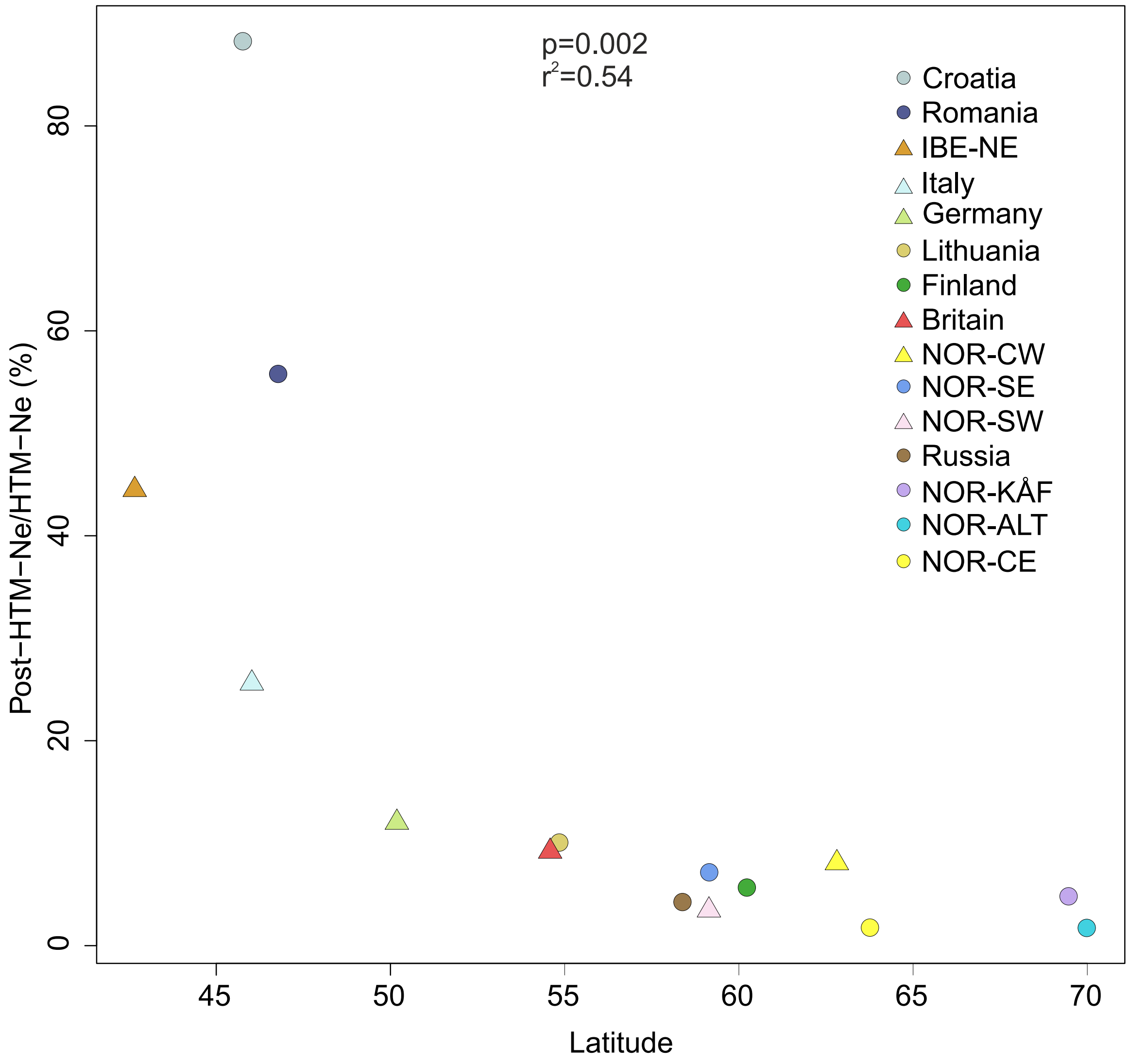


Figure S19. Decline of effective population sizes (Ne) after the Holocene thermal maximum (HTM). Y-axis shows the relative effective population size (%) after the HTM (post-HTM; 4000 ya) compared with peak values during HTM.

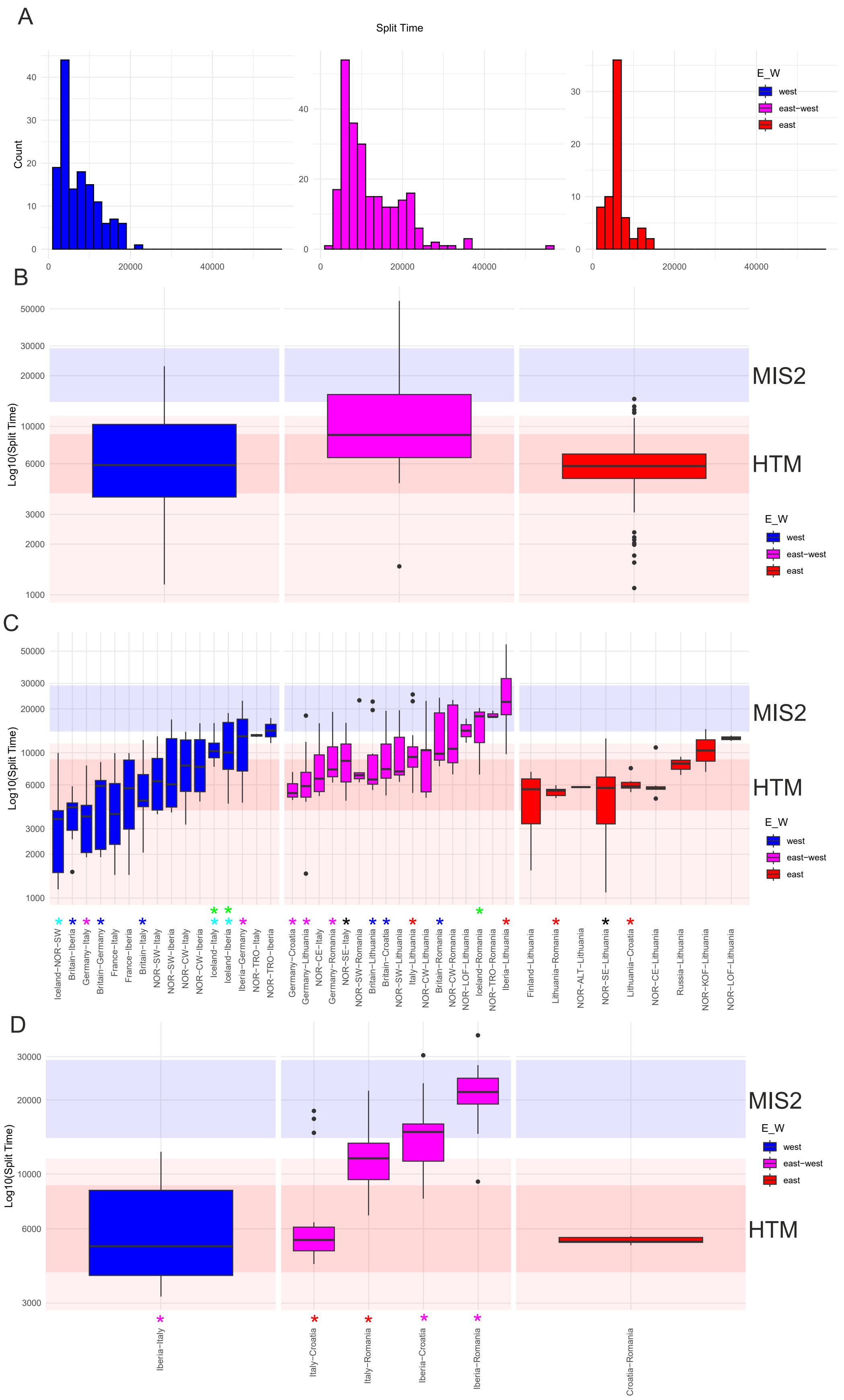
A
Split Time
Count
E_W
west
east−west
east
B
Log10(Split Time)
MIS2
HTM
C
Iceland−NOR−SW
Britain−Iberia
Germany−Italy
Britain−Germany
France−Italy
France−Iberia
Britain−Italy
NOR−SW−Italy
NOR−SW−Iberia
NOR−CW−Italy
NOR−CW−Iberia
Iceland−Italy
Iceland−Iberia
Iberia−Germany
NOR−TRO−Italy
NOR−TRO−Iberia
Germany−Croatia
Germany−Lithuania
NOR−CE−Italy
Germany−Romania
NOR−SE−Italy
NOR−SW−Romania
Britain−Lithuania
Britain−Croatia
NOR−SW−Lithuania
Italy−Lithuania
NOR−CW−Lithuania
Britain−Romania
NOR−CW−Romania
NOR−LOF−Lithuania
Iceland−Romania
NOR−TRO−Romania
Iberia−Lithuania
Finland−Lithuania
Lithuania−Romania
NOR−ALT−Lithuania
NOR−SE−Lithuania
Lithuania−Croatia
NOR−CE−Lithuania
Russia−Lithuania
NOR−KOF−Lithuania
NOR−LOF−Lithuania
D
Iberia−Italy
Italy−Croatia
Italy−Romania
Iberia−Croatia
Iberia−Romania
Croatia−Romania

Figure S20. Comparison of split times (M<0.5) between major groups (western, east-west and eastern) and subgroups (between the regions). A) Histograms of split times for major groups, B) boxplots for the same three major groups, excluding samples from southern Europe (D). C) Boxplots of split times for major groups and subgroups in north-south direction. Stars with a same color at the bottom mark significant differences in split times between western and eastern samples (Table S6) for colonization of specific central or northern regions: red (Lithuania), blue (Britain), magenta (Germany), black (Norway-SE), green (Iceland). For colonization of Iceland (cyan stars), there were also significant differences in split times within the western origin of samples. D) Split times of southern European populations (west-east direction). MIS2=Marine Isotope Stage 2 (29-14ka), Holocene (11.6-0ka) shaded by red, HTM=Holocene Thermal Maximum (9,000-4,000).

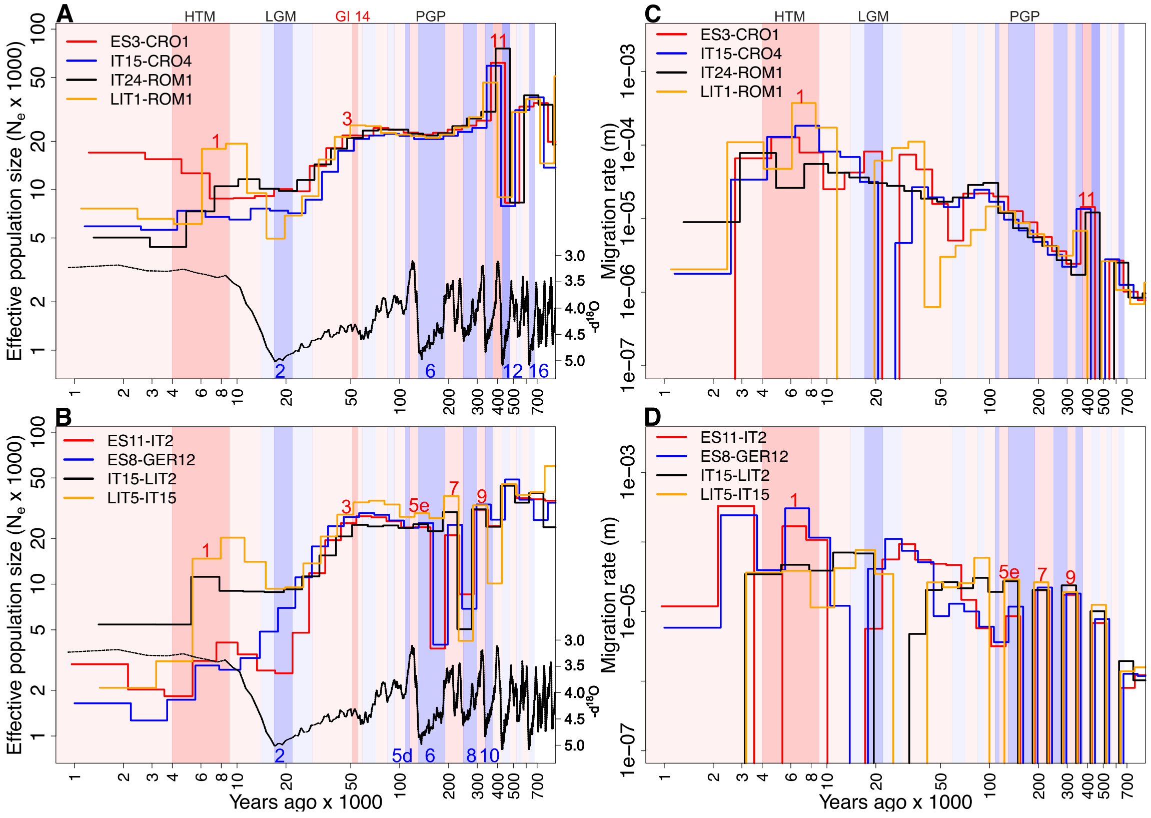


Figure S21. Climatic histories in the core and peripheral patterns with their histories extending until MIS12 and MIS10, respectively, with the acceptable resolution and cumulative migration probabilities (M<0.999). A, B) Effective population sizes ($N_E$) and migration rates of the core patterns of history (M<0.999 from MIS12 until present). C, D) Effective population sizes ($N_E$) and migration rates of the peripheral patterns of history (M<0.999 from MIS10 until present). HTM=Holocene Thermal Maximum (9,000-4,000 ya), LGM=Last Glacial Maximum (22,000-17,000 ya) GI14=Greenland Interstadial 14 (55,000-51,000 ya) PGP=Penultimate Glacial Period (190,000-130,000 ya). Curve at the bottom of A and B represents inverse benthic δ18O records from Lisiecki and Raymo (2005) and is used as a proxy for historical temperature.

**Table S1.** Genetic differentiation between regions. Weighted $F_{ST}$ between 5 samples from each region is shown.

| Region | N-KÅF | N-TRO | ICE | N-CW | N-CE | N-SW | N-SE | FIN | LIT | BRI | GER | CRO | IT | IBE-NE |
|---|---|---|---|---|---|---|---|---|---|---|---|---|---|---|
| N-ALT | 0,77 | 0,69 | 0,76 | 0,58 | 0,53 | 0,62 | 0,52 | 0,40 | 0,45 | 0,62 | 0,52 | 0,54 | 0,61 | 0,74 |
| N-KÅF | | 0,60 | 0,75 | 0,52 | 0,45 | 0,58 | 0,43 | 0,41 | 0,41 | 0,60 | 0,45 | 0,46 | 0,57 | 0,73 |
| N-TRO | | | 0,28 | 0,17 | 0,25 | 0,20 | 0,30 | 0,39 | 0,40 | 0,26 | 0,28 | 0,35 | 0,28 | 0,32 |
| ICE | | | | 0,20 | 0,28 | 0,20 | 0,34 | 0,44 | 0,45 | 0,26 | 0,31 | 0,39 | 0,31 | 0,33 |
| N-CW | | | | | 0,04 | 0,03 | 0,09 | 0,22 | 0,24 | 0,08 | 0,09 | 0,18 | 0,12 | 0,21 |
| N-CE | | | | | | 0,10 | 0,06 | 0,14 | 0,16 | 0,14 | 0,09 | 0,15 | 0,18 | 0,31 |
| N-SW | | | | | | | 0,14 | 0,28 | 0,29 | 0,11 | 0,14 | 0,23 | 0,16 | 0,26 |
| N-SE | | | | | | | | 0,11 | 0,13 | 0,18 | 0,07 | 0,13 | 0,19 | 0,36 |
| FIN | | | | | | | | | 0,05 | 0,29 | 0,15 | 0,17 | 0,29 | 0,45 |
| LIT | | | | | | | | | | 0,30 | 0,15 | 0,18 | 0,30 | 0,47 |
| BRI | | | | | | | | | | | 0,13 | 0,23 | 0,16 | 0,21 |
| GER | | | | | | | | | | | | 0,10 | 0,11 | 0,26 |
| CRO | | | | | | | | | | | | | 0,18 | 0,36 |
| IT | | | | | | | | | | | | | | 0,19 |

Abbreviations: N-ALT=Norway-Alta, N-KÅF=Norway-Kåfjord, N-TRO=Norway-Tromso, N-CW=Norway-centralwestern, N-CE=Norway-centraleastern, N-SW=Norway-southwestern, N-SE=Norway-southeastern, FIN=Finland, LIT=Lithuania, BRI=Britain, CRO=Croatia, IT=Italy, IBE-NE=Iberia-northeastern.
